# RNA promotes phase separation of glycolysis enzymes into yeast G bodies in hypoxia

**DOI:** 10.1101/638650

**Authors:** Gregory G. Fuller, Ting Han, Mallory A. Freeberg, James J. Moresco, John R Yates, John K. Kim

## Abstract

In hypoxic stress conditions, glycolysis enzymes assemble into singular cytoplasmic granules called glycolytic (G) bodies. Formation of G bodies in yeast is correlated with increased glucose consumption and cell survival. However, the physical properties and organizing principles that define G body formation are unclear. We demonstrate that glycolysis enzymes are non-canonical RNA binding proteins, sharing many common mRNA substrates that are also integral constituents of G bodies. Tethering a G body component, the beta subunit of the yeast phosphofructokinase, Pfk2, to nonspecific endoribonucleases reveals that RNA nucleates G body formation and subsequent maintenance of G body structural integrity. Consistent with a phase separation mechanism of G body formation, recruitment of glycolysis enzymes to G bodies relies on multivalent homotypic and heterotypic interactions. Furthermore, G bodies can fuse in live cells and are largely insensitive to 1,6-hexanediol treatment, consistent with a hydrogel-like state in its composition. Taken together, our results elucidate the biophysical nature of G bodies and demonstrate that RNA nucleates phase separation of the glycolysis machinery in response to hypoxic stress.

## INTRODUCTION

Cells perform many diverse activities that are spatially and temporally organized into nonmembrane bound compartments that often form transiently and can display solid, gel, or liquidlike properties. Liquid-like structures form through phase separation of protein polymers, display fast internal rearrangements, undergo fusion and fission, and exchange components with the surrounding solvent (Alberti, Gladfelter, & Mittag, 2019; Hyman, Weber, & Jülicher, 2014).

Recently, we and others demonstrated that glycolysis enzymes coalesce into membraneless cytoplasmic granules called glycolytic bodies (G bodies) in hypoxic stress in yeast, *C. elegans,* and mammalian cells (Jang et al., 2016; Jin et al., 2017; Miura et al., 2013). In yeast, the presence of G bodies correlates with accelerated glucose consumption, and impairing G body formation leads to the accumulation of upstream glycolytic metabolites (Jin et al., 2017). These data suggest that during hypoxic stress, when oxidative phosphorylation is inhibited, G bodies form to enhance the rate of glycolysis by concentrating glycolysis enzymes. In *C. elegans,* hypoxia rapidly induces the formation of foci containing glycolysis enzymes near presynaptic release sites in neurons. A phosphofructokinase mutant incapable of punctate localization disrupts synaptic vesicle clustering in neurons, suggesting that coalescence of glycolysis enzymes promotes synaptic function (Jang et al., 2016). However, the mechanism of G body formation remains poorly understood.

G bodies have a number of features common to known phase-separated bodies. In addition to being non-membrane bound, some G body components, including phosphofructokinase 2 (Pfk2p), contain intrinsically disordered regions (IDRs), a feature of proteins that undergo phase transitions (Jin et al., 2017). The IDR is required for Pfk2p localization to G bodies (Jin et al., 2017). In addition, G body formation in mammalian cells is inhibited by addition of RNase to the culture media, suggesting that RNA is required for G body integrity (Jin et al. 2017).

Many other phase-separated structures contain RNA. For instance, stress granules contain mRNAs stalled in translation initiation (Buchan, Muhlrad, & Parker, 2008) and involve protein-protein and IDR interactions between mRNA binding proteins (Panas, Ivanov, & Anderson, 2016; Protter et al., 2018). Nucleoli, which are sites of ribosomal RNA processing, also form by phase separation (Brangwynne, Mitchison, & Hyman, 2011; Protter et al., 2018).

For some proteins, RNAs can promote phase separation *in vitro* (Elbaum-Garfinkle et al., 2015; Zhang et al., 2015) or even phase separate by themselves (A. Jain & Vale, 2017; Van Treeck et al., 2018). Furthermore, RNase treatment can disrupt mature granules such as P bodies, demonstrating the importance of RNAs in the structural integrity of RNP granules (Teixeira, Sheth, Valencia-Sanchez, Brengues, & Parker, 2005). The identity of bound RNAs can be important for phase separation. For example, the yeast Whi3p protein phase separates in the presence of its substrate RNA, *CLN3,* but not in the presence of total RNA (Zhang et al., 2015). Additionally, RNAs that phase separate alone from yeast total RNA *in vitro* are enriched in stress granules, suggesting that RNA may drive phase separation of stress granule proteins *in vivo* (Van Treeck et al., 2018).

In contrast, other RNA binding proteins display the opposite behavior. For Pab1, a core stress granule component, high concentrations of RNA prevent phase separation *in vitro* (Riback et al., 2017). Microinjection of RNase A can induce aggregation of nuclear FUS protein *in vivo,* suggesting that, in this case, high levels of RNA oppose phase separation. RNA binding-defective mutants of TDP-43 display an increased propensity to phase separate *in vitro* and *in vivo* and addition of TDP-43 RNA substrates allows TDP-43 to remain soluble (Maharana et al., 2018; J. R. Mann et al., 2019).

Taken together, reconciling these disparate behaviors in which some RNAs promote formation of stress granules and P bodies, while others antagonize phase separation of proteins such as TDP-43 and FUS will require targeted *in vivo* approaches to determine how mature granules are affected by RNA. In granules for which RNA facilitates RNP granule formation, RNA may function by forming a scaffold to promote multivalent interactions (Fay & Anderson, 2018; A. Jain & Vale, 2017; Langdon & Gladfelter, 2018). Interestingly, metabolic enzymes, including some involved in glycolysis, bind mRNAs (Beckmann et al., 2015; Matia-González, Laing, & Gerber, 2015). However, the physiological role(s) of this RNA binding remain poorly understood.

In this study, we show that G bodies are novel RNP granules formed through a phase separation mechanism *in vivo.* We identify the common mRNA substrates of the G body resident glycolysis machinery and demonstrate the essential role that RNA plays in G body biogenesis and maintenance *in vivo.* Thus, our data suggest a model where, in response to hypoxic stress, when cellular demand for energy must be met solely through glycolysis, G bodies form through multivalent protein-protein and protein-RNA interactions to enhance the rate of glycolysis.

## RESULTS

### Analysis of the RNA-binding proteome uncovers core G body constituents

To identify the RNA binding proteome in S. *cerevisae,* we incorporated 4-thiouridine (4Su) into RNA in log phase yeast cells grown under normoxic conditions and combined photoactivatable-ribonucleoside-enhanced cross-linking (PAR-CL) with oligo(dT) affinity purification and tandem mass spectrometry (PAR-CL-MS, Fig 1A) as previously described (Baltz et al., 2012; Castello et al., 2012). Stringent washes and separation by SDS-PAGE removed non-crosslinked proteins. We identified 259 mRNA-binding proteins (mRBPs), nearly half of which were non-canonical RBPs that do not contain conventional RNA binding domains. Many of these have been identified as mRBPs in yeast and other organisms (Beckmann et al., n.d.; Matia-González et al., 2015), thus validating our approach (Fig 1B). In addition, our study identified 69 novel mRBPs in yeast (Fig 1B, Table S1). Consistent with previous studies, we identified 10 glycolysis enzymes as mRBPs and other studies identified 5 additional glycolysis enzymes (Beckmann et al., 2015; Matia-González et al., 2015). Of the RNA binding glycolysis enzymes 10 localize to G bodies in hypoxia (Fig S1A), (Jin et al., 2017).

**Figure 1:**
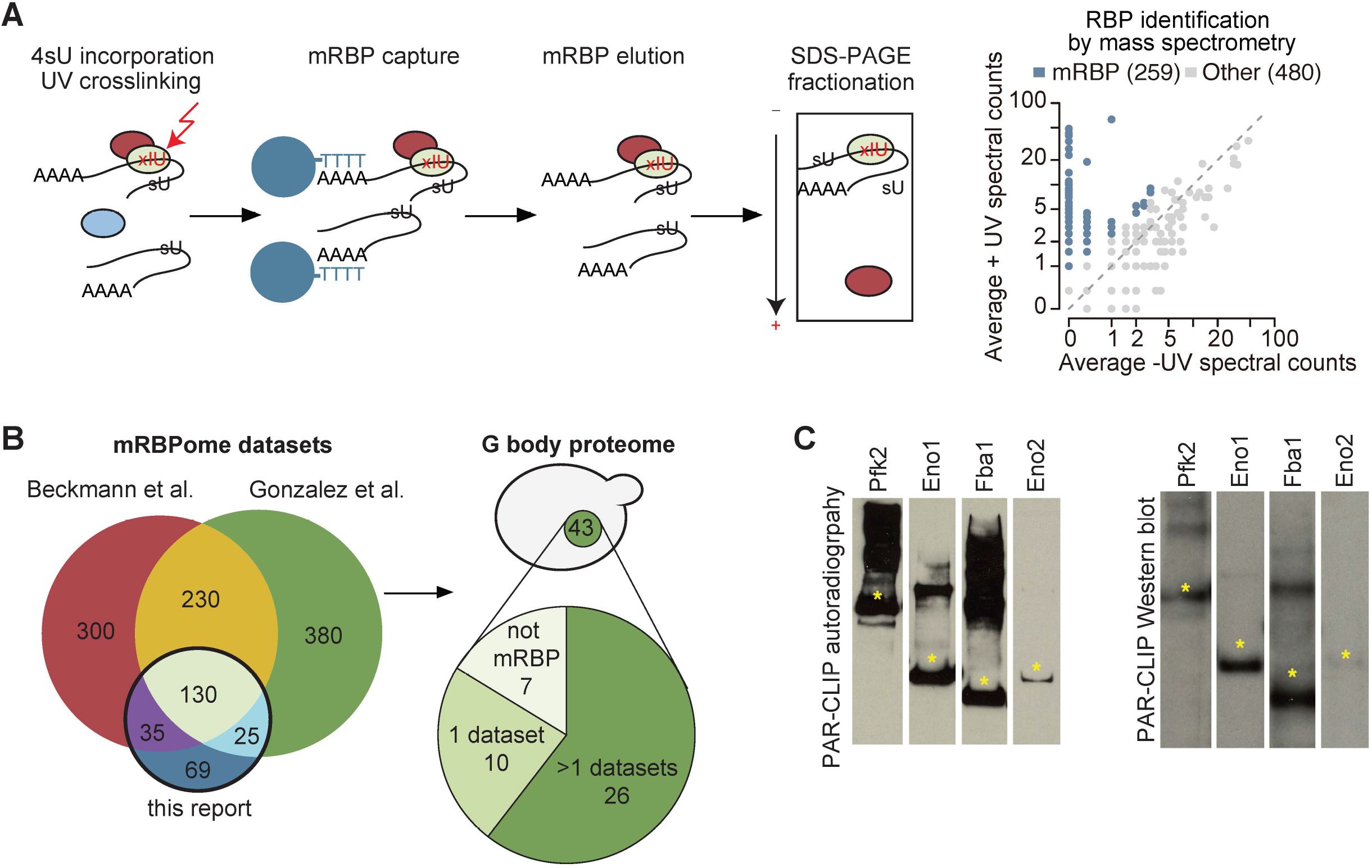
G bodies are enriched for RNA-binding proteins. (A) PAR-CL-MS pipeline. 4 Thiouridine (4Su) is incorporated by supplementation in media and crosslinked to protein with 365 nm UV light. mRBPs are captured with oligo-d(T) beads and eluted. RNPs are fractionated by SDS-PAGE and resulting proteins are analyzed by proteomic mass spectrometry. Average mass spectrometry spectral peak counts from biological replicate PAR-CL-MS versus non-UV-treated control. Proteins enriched in PAR-CL-MS (blue dots) had either (1) >2 spectral counts in PAR-CL and 0 spectral counts in -UV control or (2) had FDR <10% as calculated by QSPEC (Choi, Fermin, & Nesvizhskii, 2008). (B) Overlap of identified mRBPs with datasets generated with the same methodology. mRBP datasets are from (Matia-González et al., 2015); (Beckmann et al., 2015). Distribution of G-body proteome (union of colocalization validated G-body proteins from (Jin et al., 2017)(Miura et al., 2013). (C) Autoradiography and western blot of PAR-CLIP of TAP-tagged proteins (Pfk2, Eno1, Eno2, and Fba1). Stars indicate the same band in autoradiography and western blot confirming RNA binding by each enzyme.

### Glycolysis enzymes bind similar transcripts

To validate RNA binding by glycolysis enzymes, we end-labeled potential RNAs crosslinked to TAP-tagged Pfk2, Eno1, Eno2, and Fba1 (Fig 1C, left panels) with γ-32-P ATP. The autoradiographs displayed the same migration pattern in SDS-PAGE as the immunoblots of the corresponding TAP-tagged glycolysis enzymes (Fig. 1C, right panels), suggesting that the glycolysis enzymes Pfk2, Eno1, Eno2, and Fba1 bind RNA. We then identified the mRNA substrates of Pfk2, Eno1, and Fba1 by performing photoactivatable-ribonucleoside-enhanced cross-linking and immunoprecipitation (PAR-CLIP) followed by deep sequencing (PAR-CLIP-seq; (Hafner et al., 2010) in log phase yeast cells grown under normoxic conditions. To determine the binding site sequences, we empirically determined “h¡gh-conf¡dence” RPM thresholds (Pfk2: 5 RPM, Fba1: 0.5 RPM, Eno1: 0.5 RPM) for each library (Fig S1B). We identified 1,540 total mRNAs that bind at least one of these three glycolysis enzymes. Specifically, there were 439 direct mRNA substrates of Pfk2 with 559 discrete binding sites, 1,001 mRNA substrates with 1,432 binding sites for Eno1, and 721 mRNA substrates with 1,014 binding sites for Fba1 (Fig S1B, right panels). The results of Eno1 PAR-CLIP-seq were in agreement with a recent analysis of Eno1 in normoxic conditions by CRAC (a method that UV crosslinks and affinity purifies protein-RNA complexes under denaturing conditions (Shchepachev et al., 2019)), providing additional validation of our identified binding sites. We identified 69 out of the top 100 bound mRNAs in the CRAC dataset. mRNAs bound by each glycolysis enzyme displayed substantial overlap, with 490 mRNAs binding at least two of the three glycolysis enzymes and 131 mRNAs binding all three (Fig 2A). Intriguingly, bound mRNAs of all three enzymes were enriched for functional annotations related to glycolysis as well as other metabolic pathways (Fig 2B). For example, Pfk2 targets included mRNAs encoding 13 of the 22 known glycolytic enzymes (Fig S1C). Fba1 and Eno1 also bound to mRNAs encoding glycolysis enzymes, and together, Pfk2, Fba1, and Eno1 bound to 16 of the 22 glycolytic enzyme-encoding mRNAs in yeast (Fig S1C).

**Figure 2:**
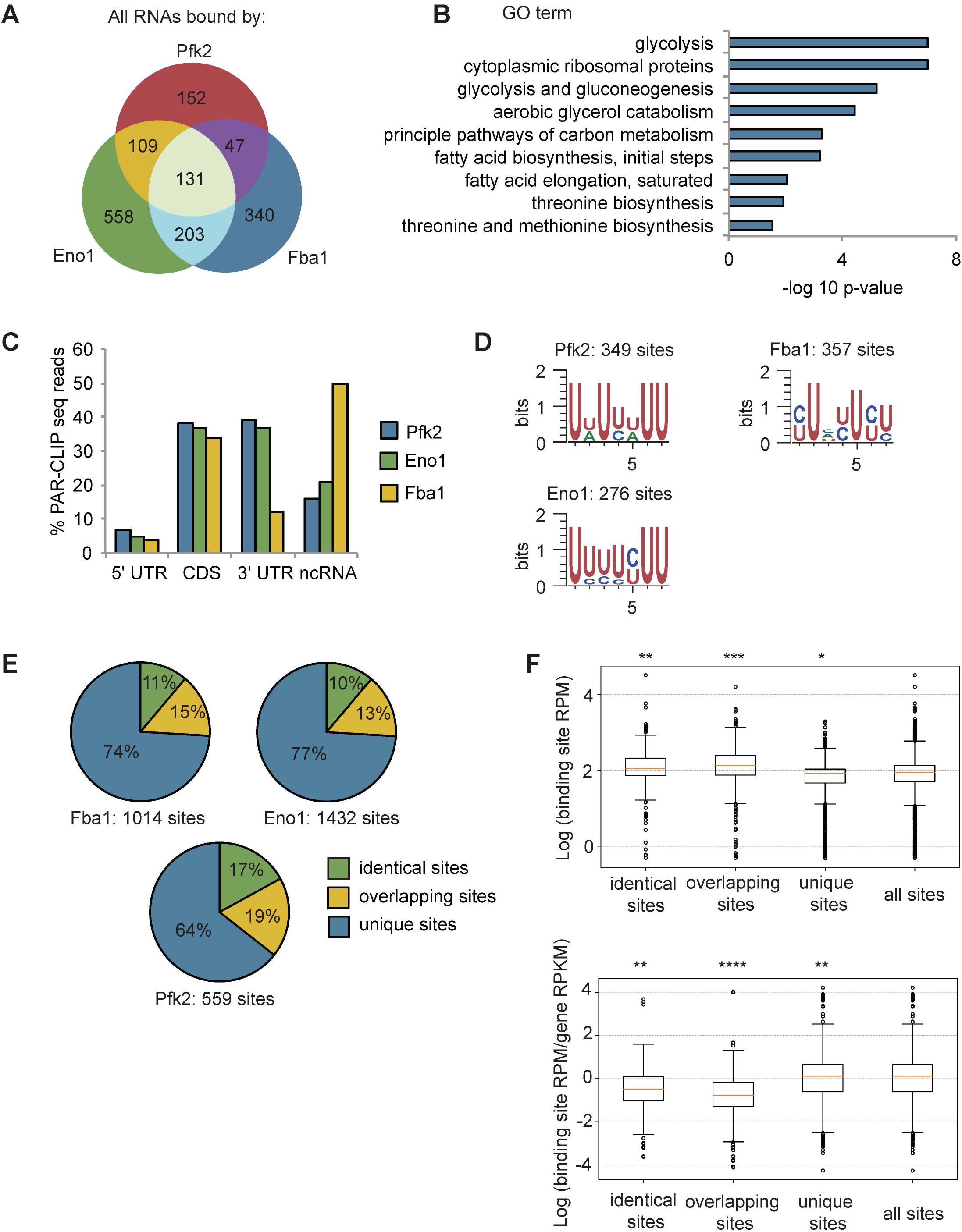
Glycolysis enzymes bind similar RNAs. (A) Overlap of mRNAs bound by Pfk2, Eno1, and Fba1 as identified by PAR-CLIP Seq in normoxic conditions. B. (B) Gene ontology (GO) terms enriched among transcripts containing high-confidence Pfk2, Eno1 and Fba1 sites include glycolysis. Fisher’s exact test p-values are plotted. (C) Percent of total Pfk2, Eno1, and Fba1 PAR-CLIP-seq reads per million mapped reads (RPM), aligning with the indicated genic regions. (D) Identified sequence motifs among Pfk2, Eno1, and Fba1 binding sites. (E) Percent of binding sites for each PAR-CLIP Seq dataset that are bound by more than one glycolysis enzyme (identical sites), overlap a binding site of another glycolysis enzyme (overlapping sites), or are bound by only one glycolysis enzymes (unique sites) (F) (Top) Binding sites present in multiple datasets or overlapping other binding sites tend to have greater read depth. Boxplot of Log_10_ (Binding Site RPM) for each class of binding site. p-values represent unpaired student’s T Tests. (Bottom) Glycolysis enzymes tend to bind tighter to unique sites. Boxplot of Log_10_ ratio of binding site RPM to gene RPKM from mRNA seq. p-values represent unpaired student’s T tests. * P<10^-5^. ** P<10^-7^. *** P<10^-9^. ****P<10^-20^.

Most of the glycolytic enzyme binding sites on substrate mRNAs were within the 3’ untranslated regions (3’ UTRs) and coding sequences, followed by non-coding RNAs and 5’ UTRs (Fig 2C). We used the high-confidence binding sites for each glycolysis enzyme to identify enriched motifs. Pfk2 binding sites contained an AU-rich element, similar to elements that regulate mRNA stability in yeast (Vasudevan & Peltz, 2001), whereas Eno1 and Fba1 binding sites contained pyrimidine-rich motifs (Fig 2D). Furthermore, the binding sites for each enzyme displayed overlap between enzymes. Although binding footprints were short (22 nt on average), between 10%-17% of binding sites were bound by at least two of the three glycolysis enzymes (Fba1, Eno1, or Pfk2), and 13%-19% of the sites for one enzyme partially overlapped with the binding sites of at least one other glycolysis enzyme with the remainder of binding sites being uniquely bound by either Pfk2, Eno1 or Fba1 (Fig 2E). These partially overlapping sites and identical shared sites had greater average sequencing depth than unique sites bound by only one glycolysis enzyme (Fig 2F). Unique binding sites, however, had a greater log enrichment of binding site RPM to gene reads per kilobase million (RPKM) than overlapping sites and identical sites in multiple datasets, suggesting that these sites were more tightly bound (Fig 2F). Thus, the greater binding frequency on overlapping sites was largely driven by the amount of target mRNA. Nevertheless, the overlap in bound transcripts and binding sites suggests that common RNA binding could contribute to the coalescence of glycolysis enzymes into G bodies.

### G bodies copurify with RNAs

We previously developed a method using differential centrifugation and affinity capture to isolate G bodies and identify their resident proteins by mass spectrometry followed by validation using colocalization to G bodies (Jin et al., 2017). Of the 43 identified G body components, 36 are mRBPs (Jin et al., 2017; Miura et al., 2013) (Fig 1B, Table S1). The observation that many of the proteins targeted to G bodies upon shift to hypoxic conditions, including glycolysis enzymes, bind RNA raised the possibility that G bodies themselves contain RNA. To test this hypothesis, we adapted our previous G body purification protocol to perform RNA extraction, cDNA synthesis, and quantitative PCR to evaluate the presence of co-purifying RNA (Fig S2A). In order to reduce background, we generated a yeast strain with an integrated Pfk2-GFP-1xFlag transgene. This yeast strain was shifted to hypoxic conditions for 18 hours to induce G body formation. G bodies were then immunopurified from lysates with a monoclonal mouse anti-Flag antibody and eluted with Flag peptide before proteinase K digestion and RNA extraction (Fig S2A-B). As a control for nonspecific RNA binding, we performed the same protocol with wild type BY4742 cells and extracted RNA from the flow through for each sample (Fig S2A-B). By comparing the relative amount of RNA from the flow through and the eluate, we determined the percent of RNA in the eluate compared to the input of the immunoprecipitation. We tested a range of qPCR probes for RNAs with or without binding sites identified in PAR-CLIP-seq of Eno1p, Pfk2p and Fba1p (Table S2). While Pfk2-GFP-Flag pulldowns recovered between 2.4% and 7% of the flow-through RNA, we recovered at most 1.5% of flow-through RNA in control experiments (Fig S2C), indicating a 3.5 to 11.5-fold enrichment of these RNAs in G bodies versus the BY4742 control. These trends were not different for RNAs with binding sites for glycolysis enzymes, suggesting that RNA binding by other RBPs in G bodies contribute to RNA accumulation in G bodies. However, the low recovery rate relative to flow-through RNAs suggests that only a small fraction of each mRNA accumulates in G bodies, unlike stress granules, where up to 95% of particular mRNAs localize to stress granules (Khong et al., 2017).

### Targeting RNase to nascent G body sites inhibits formation *in vivo*

Since G bodies are enriched for RNA binding proteins and because G body resident glycolysis enzymes share many common mRNA substrates, we next tested whether G body formation requires RNA *in vivo.* We targeted an RNAse to nascent G bodies by fusing Pfk2 to *E. coli* MqsR, an RNase that preferentially cleaves RNA at GCU sequences, followed by a C-terminal Flag tag (Kasari, Kurg, Margus, Tenson, & Kaldalu, 2010; Yamaguchi, Park, & Inouye, 2009). We placed Pfk2p-MqsR-Flag under the control of the copper sulfate (CuSO_4_) inducible *CUP1* promoter on a centromeric plasmid, and introduced this plasmid, or a control vector, into cells expressing the G body reporter Pfk2-GFP (Fig 3A). We detected weak expression of Pfk2-MqsR-Flag even in the absence of CuSO_4_, consistent with weak activation of the *CUP1* promoter in hypoxic conditions (Becerra et al., 2002). Cells treated with CuSO_4_ showed a dose-dependent increase in Pfk2-MqsR-Flag levels (Fig S3A). At low concentrations of CuSO_4_ (5, 10 μM) in hypoxia, the Pfk2-MqsR-Flag fusion protein was targeted to G bodies and colocalized with Pfk2-GFP (Fig S3B-C). Using this same approach, another RNase, RNaseA, could also be induced and targeted to G bodies as a Pfk2-RNase A fusion protein (Fig S3D, E).

**Figure 3:**
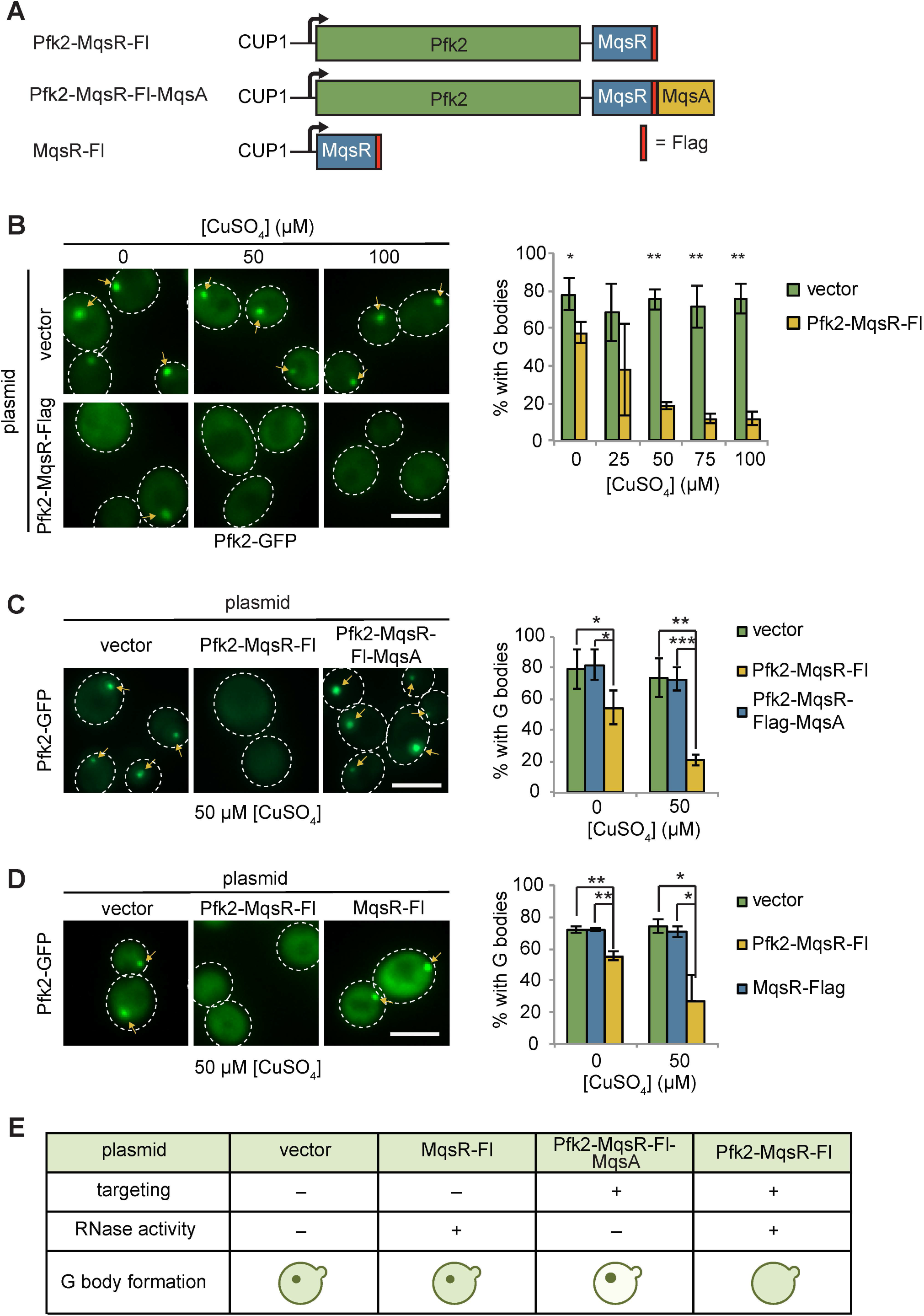
Tethering Pfk2 to an RNase prevents G-body formation. (A) Schematic of Pfk2-MqsR constructs. All constructs were expressed from a centromeric plasmid under control of the copper sulfate inducible CUP1 promoter. (B) (Left) Representative images of hypoxic Pfk2-GFP localization with increasing concentrations of copper sulfate for cells expressing a vector control or cells expressing Pfk2-MqsR-Flag. (Right) Quantification of G body formation in cells with varying concentration of CuSO_4_. (C) (Left) Representative images of hypoxic Pfk2-GFP localization comparing cells with a vector control, Pfk2-MqsR-Fl, or Pfk2-MqsR-Fl-MqsA with 50 μM CuSO_4_. (Right) Quantification of G body formation for cells with each plasmid with 0 and 50 μM CuSO_4_. (D) (Left) Representative images of hypoxic Pfk2-GFP localization at 50 μM CuSO_4_. (Right) Quantification of G body formation for cells with each plasmid with 0 and 50 μM CuSO_4_. (E) Cartoon and summary of results of G body formation with either a vector plasmid or plasmids expressing either MqsR-Flag, Pfk2-MqsR-Flag, or Pfk2-MqsR-Flag-MqsA. All scale bars are 5 μM. For each graph, data represent mean and standard deviation of three to four individual experiments (N > 100 per replicate per condition). Arrows indicate G bodies. Statistics were analyzed by unpaired student’s T tests with a Bonferroni correction for multiple testing. * P<0.05. ** P<0.01. *** P<0.001

Targeting Pfk2-MqsR-Flag to G bodies caused a robust reduction in the fraction of hypoxic cells with G bodies in a dose-dependent manner (Fig 3B). In contrast, G body formation in hypoxic cells carrying a control vector was unaffected by CuSO_4_ treatment at all concentrations tested (Fig 3B). Even without induction, hypoxic cells carrying the Pfk2-MqsR-Flag plasmid showed a 20% decrease in G body formation, consistent with the weak expression of Pfk2-MqsR-Flag under these conditions. CuSO_4_-induced expression of Pfk2-RNase A similarly inhibited G body formation (Fig S4A).

The MqsR RNase is inhibited by its antitoxin, MqsA (Kasari et al., 2010; Yamaguchi et al., 2009). To verify that MqsR nuclease activity was required to inhibit G body formation, we fused the antitoxin MqsA to the Pfk2-MqsR-Flag construct, generating a Pfk2-MqsR-Flag-MqsA fusion protein. At both 0 and 50 μM CuSO_4_, G body formation in hypoxic cells was now unaffected by induction of Pfk2-MqsR-Flag-MqsA (Fig 3C), indicating that MqsA effectively inhibited the G body-targeted MqsR RNase from disrupting G body formation. G bodies appeared larger and brighter in Pfk2-MqsR-Flag-MqsA-expressing cells than in cell expressing a vector control, possibly due to overexpression of Pfk2 (Fig. 3C). Similarly, a Pfk2-RNase A^H12A^ mutant with decreased RNase activity was less effective at reducing G body formation than Pfk2-RNase A^wild-type^, especially at 100 μM CuSO_4_ (Fig S4B). Pfk2-RNase A^H12A^ possesses residual RNase activity (Thompson & Raines, 1994), which likely contributes to the modest loss of G body formation. Thus, inhibition of G body formation by RNase A and MqsR is due to their ribonuclease activity.

To further test whether specifically targeting of MqsR-Flag to G bodies by fusion to Pfk2 was required to inhibit G body formation, we engineered cells expressing MqsR-Flag alone, which was inducible by addition of CuSO_4_ (Fig S3A). Unlike Pfk2-MqsR-Flag expression, MqsR-Flag expression had no effect on G body formation in hypoxic cells (Fig 3D). In addition, we tested the ability of Pfk2-RNase A to inhibit G body formation of several other G body markers, including Eno2, Cdc19, and Fba1. In each case, CuSO_4_ induction of hypoxic cells expressing Pfk2-RNase A, but not the vector control, led to a decrease in G body formation (Fig S5).

### Targeting RNase to pre-formed G bodies causes multiple puncta formation

To test whether RNA was required for the stability of G bodies that had already formed, cells were first grown in hypoxic conditions for 20 h and then shifted to normoxic conditions along with Pfk2-MqsR-Flag induction by addition of CuSO_4_. Immediately following shift from hypoxic to normoxic conditions and concomitant CuSO_4_ addition (i.e., “0 h post-hypoxia” in Fig 4A, B), Pfk2-MqsR-Flag carrying cells showed a modest reduction in G body formation compared to controls (Fig 4A, B), likely due to leaky expression of Pfk2-MqsR-Flag in hypoxia. However, we reasoned that we could still test the effects of inducing Pfk2-MqsR, as most cells (61%) had G bodies. After prolonged induction of Pfk2-MqsR-Flag in normoxic conditions (“20 h post-hypoxia” in Fig 4D, E), we observed strong expression of Pfk2-MqsR-Flag at 100 μM CuSO_4_ and low-level expression even in cells without addition of CuSO_4_ (Fig 4C). Under these conditions, cells harboring a vector control either had a single G body per cell that persisted or diffuse localization of Pfk2-GFP, regardless of added CuSO_4_. In contrast, a substantial proportion of cells (41-50%) with Pfk2-MqsR-Flag showed multiple puncta per cell (Fig. 4D-E). These data suggest that pre-formed G bodies fracture into multiple structures in the presence of Pfk2-MqsR-Flag.

**Figure 4:**
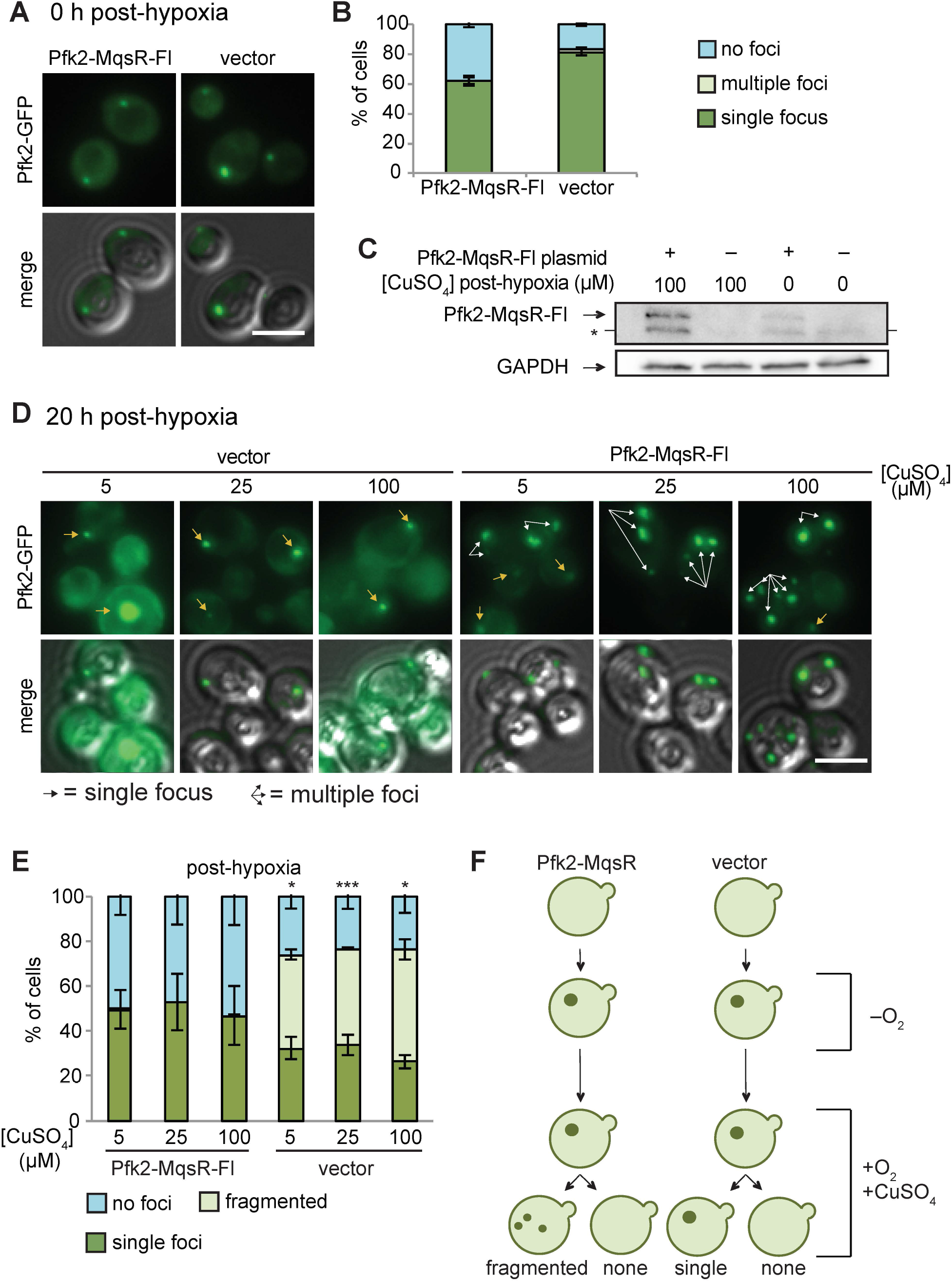
Pfk2-MqsR-Fl induction fractures existing G bodies. (A) Cells with a plasmid inducibly expressing Pfk2-MqsR-Flag form G bodies in the absence of induction by CuSO_4_. Representative images of hypoxic Pfk2-GFP localization in cells expressing a vector control or Pfk2-MqsR-Fl with 0 μM CuSO_4_. (B) Quantification of G-body formation of cells in (A). (C) Western blot showing induction of Pfk2-MqsR-Fl where CuSO_4_ is added after 20 h in hypoxia and cells are subsequently cultured for 20 h in normoxia. Pfk2-MqsR-Fl is probed with a monoclonal anti-flag antibody. GAPDH serves as a loading control. * indicates a nonspecific band. (D) Representative images of Pfk2-GFP localization in cells expressing a vector control and cells expressing Pfk2-MqsR-Fl after 20 h hypoxia followed by induction with varying concentrations of CuSO_4_ in normoxia for 20 h. Cells with Pfk2-MqsR-Fl frequently have multiple large foci. (E) Quantification of cells with a single focus, multiple foci, or no foci for cells in (D). (F) Cartoon showing Pfk2-GFP localization before hypoxia, immediately after hypoxia, and after induction with CuSO_4_. All graphs show mean and standard deviation of three independent experiments (N > 100 cells per condition per replicate). Arrows indicate G bodies. Statistics calculated with unpaired student’s T tests. All scale bars 5 μm. * P<0.05. ** P<0.01. *** P<0.001

Taken together, these data indicate that 1) Nascent G body formation was inhibited by RNases in a concentration-dependent manner (Fig. 3A); 2) Inhibition of de novo G body formation required both RNase activity and targeting to the site of G body formation (Fig. 3C-E); 3) Once formed, targeting an RNase to G bodies led to cells with multiple foci, indicating disruption but not dissolution of G bodies (Fig. 4F); and 4) Once formed in hypoxia, G bodies could persist for tens of hours in normoxia (Fig. 4D).

### G body recruitment requires multivalent interactions

IDRs have been shown to modify phase separation behavior *in vitro* and *in vivo* (Kato et al., 2012; Mitrea & Kriwacki, 2016; Molliex et al., 2015; Protter et al., 2018). Pfk2 has such an IDR in its N-terminal domain (Fig 5A). Our previous study demonstrated that deletion of an N-terminal IDR spanning amino acids 140-165 of Pfk2-GFP (Pfk2^Δ140-165^-GFP) increased the number of cells forming multiple foci and decreased overall G body formation (Jin et al., 2017). The Pfk2 N-terminal domain is conserved in S. *cerevisiae* (28.7% identity) and other yeast species. The crystal structure of S. *cerevisiae* heterooctameric phosphofructokinase (with four copies each of Pfk1 and Pfk2) lacks this region for both Pfk1 and Pfk2, due to proteolytic cleavage in sample preparation (Banaszak et al., 2011). However, the crystal structure of phosphofructokinase from a related yeast species, *Komegataella pastoris* (Strater et al., 2010), shows that the Pfk2 N-terminal domain resembles a glyoxylase domain and interacts with Pfk1. Regions outside of the Pfk2 N-terminus also interact with Pfk1 in both crystal structures to form a heterooctamer with four Pfk2 and four Pfk1 molecules. These multiple interaction domains provide the potential for multivalent Pfk2 interactions.

**Figure 5:**
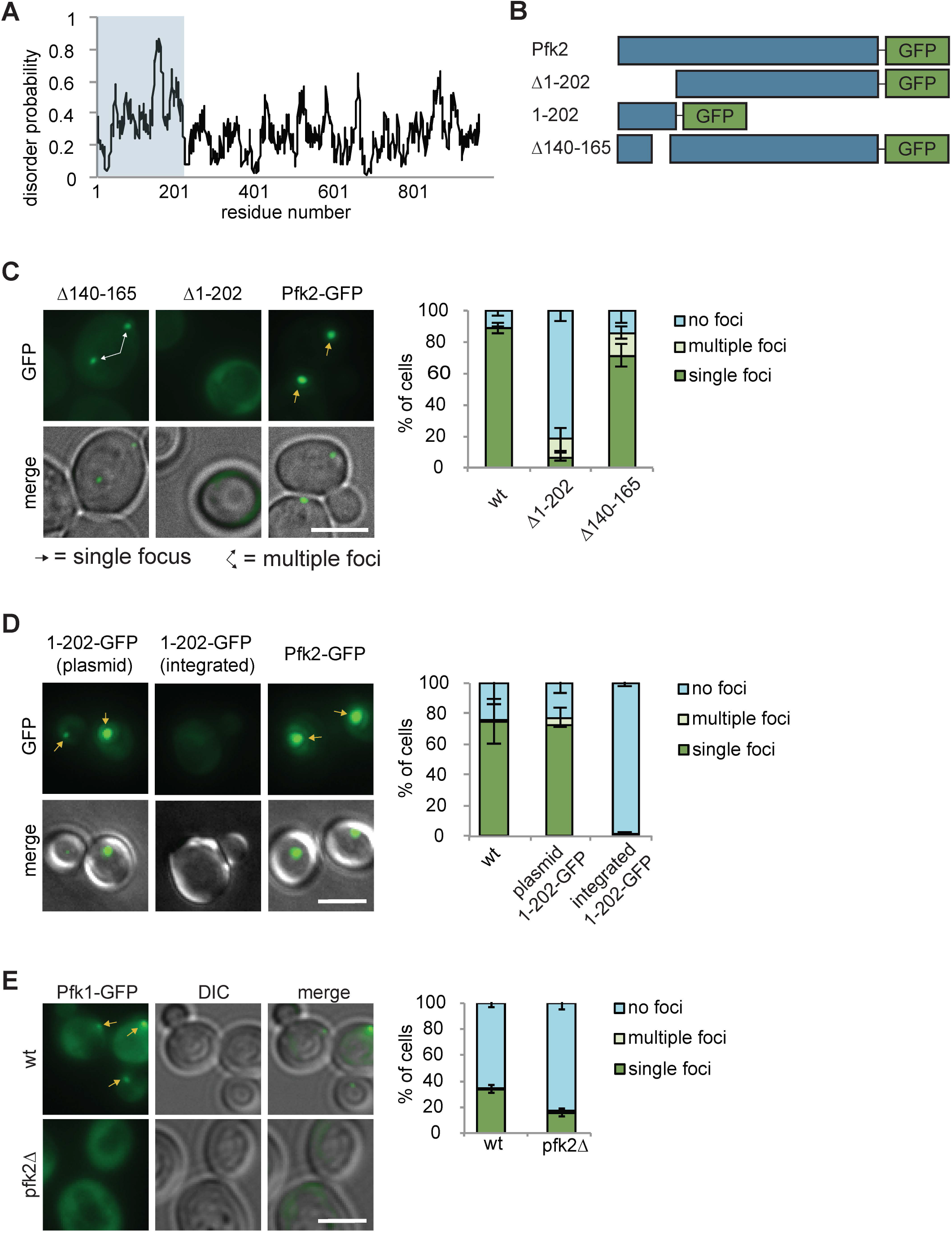
Protein recruitment to G bodies relies on multivalent interactions. (A) IUPRED prediction of disorder in Pfk2. The first 202 residues are largely unstructured. (B) Schematic of Pfk2-GFP variants tested. (C) (Left) The Pfk2 N terminus is required for G-body localization. Representative images of Pfk2-GFP variant localization in hypoxia. (Right) Quantification of G-body localization. (D) (Left) The Pfk2 N terminus is not sufficient for G-body localization. Representative images of Pfk2-GFP variant localization in hypoxia. When integrated (in the absence of full-length Pfk2), the Pfk2 N terminus does not localize to G bodies. When expressed on a plasmid in a strain with full-length Pfk2, the Pfk2 N terminus localizes to G bodies. (Right) Quantification of G-body localization. (E) (Left) Pfk1-GFP recruitment to G bodies depends on Pfk2. Representative images of hypoxic Pfk1-GFP localization in wild-type and *pfk2Δ* cells. (Right) Quantification of Pfk1-GFP localization in wild-type and *pfk2Δ* cells in hypoxia. All scale bars are 5 μm. Arrows represent either G bodies or cells with multiple G bodies. Data represent mean and standard deviation of three replicates (N > 100 cells per condition per replicate). Statistics are unpaired student’s T tests comparing G-body formation. * P<0.05. ** P<0.01. *** P<0.001

To test whether N-terminal interactions are important for Pfk2 recruitment to G bodies, we deleted its N-terminal domain (Δ1-202) (Fig 5B). This region encompasses the most disordered residues including the previous IDR deletion (Δ140-165) (Fig 5B). Compared with full-length Pfk2-GFP, cells expressing Pfk2^Δ1-202^-GFP had drastically reduced puncta (Fig 5C), suggesting that the N-terminal domain is required for either G body formation or Pfk2 localization to G bodies. As a control, we retested the Pfk2^Δ140-165^-GFP mutant and observed decreased G body localization with an increased instance of cells with multiple foci. Notably, the localization defect was much stronger in Pfk2^Δ1-202^-GFP cells suggesting that the structured portions of the N-terminal domain act in concert with the disordered residues to promote localization to G bodies.

To test whether the N-terminal region is sufficient for G body localization, we fused the N-terminal region alone to GFP (Pfk2^1-202^-GFP, Fig 5B). When *pfk2^1-202^-gfp* was integrated and replaced full-length endogenous *pfk2,* Pfk2^1-202^-GFP did not form puncta (Fig 5D). However, when Pfk2^1-202^-GFP was expressed from a plasmid in wild-type cells expressing endogenous full-length Pfk2, we detected robust puncta formation (Fig 5D). These data suggest that the N-terminal region of Pfk2 is not sufficient for G body localization, but requires interaction with full-length Pfk2 for recruitment to G bodies. To test the importance of the ordered interactions by the remainder of the protein, we probed Pfk1 recruitment to G bodies in wild-type and *pfk2Δ* cells. The frequency of hypoxic cells with Pfk1-GFP puncta was reduced in *pfk2Δ* cells compared to wild-type, suggesting that Pfk1 is recruited to G bodies in concert with Pfk2 (Fig 5E).

### G bodies behave as gels

To determine if G bodies behave as liquid-like structures or more solid gels, we treated hypoxic cells with 5% 1,6-hexanediol. The alcohol 1,6-hexanediol specifically dissolves liquid-like structures purportedly due to its ability to disrupt weak hydrophobic interactions, and therefore has been used to discern liquid-like from solid-like granules (Kroschwald, Maharana, & Simon, 2017; Kroschwald et al., 2015). We noticed no significant loss of G bodies after 1 h, whereas untreated cells showed a modest loss of G bodies (Fig S6A). The loss of G bodies in the control condition was likely due to continued cell division, which is inhibited by 1,6-hexanediol (Kroschwald et al., 2017). However, G bodies in 1,6-hexanediol treated cells appeared smaller than those in untreated cells. Therefore, we quantified the size of G bodies by fitting 2-dimensional Gaussian distributions to maximum intensity projections of G bodies in untreated and 1,6-hexanediol treated cells and computed the parameters of the fit. In 1,6-hexanediol treated cells, the average σ_X_ (0.72 +/− 0.055 μm) and σ_Y_ (0.74 +/− 0.060 μm) values were lower than untreated cells (1.1 +/− 0.070 μm and 1.1 +/− 0.067 μm respectively) indicating a smaller granule radius (Fig S6A). Additionally, the amplitude was lower in the 1,6-hexanediol treated cells (12,054 +/− 857 A.U. compared to 18,519 +/− 1308 A.U.), indicating reduced total fluorescence in the granules. Thus, while 1,6-hexanediol does not fully dissolve G bodies, it can reduce their size. As a control, we verified that 1,6-hexanediol treatment of glucose-starved cells disrupted P bodies labeled with Edc3-GFP, but caused only a small and not significant loss of stress granules labeled with Pab1-GFP (Kroschwald et al., 2015) (Fig S6B). To test whether an early liquid-liquid phase separation (LLPS) state precedes G body formation, we treated cells with 1,6-hexanediol during nascent granule formation. Because high concentrations (5%) of 1,6-hexanediol severely inhibited cell growth, we tested G body formation with 2% 1,6-hexanediol. As a control, we verified that addition of 1,6-hexanediol to the starvation media inhibited the formation of P bodies but not stress granules (Fig S6C) (Kroschwald et al., 2017). We found that 1,6-hexanediol treatment of Pfk2-GFP wild-type cells during growth in hypoxia did not affect nascent G body formation (Fig S6D), indicating that 1,6-hexanediol cannot prevent nucleation of G bodies.

To test if a mutant with decreased G body formation would be more susceptible to disruption by 1,6-hexanediol, we used *snf1Δ* cells. We previously demonstrated that cells lacking Snf1, the yeast ortholog of AMP activated protein kinase, had decreased G body formation and frequently formed multiple smaller foci per cell, rather than the single foci observed in wild type cells (Jin et al., 2017). Unexpectedly, 1,6-hexanediol treatment of *snf1Δ* cells increased the frequency of cells with a single Pfk2-GFP labeled G body (Fig. S6D), such that they appeared superficially wild type. Therefore, 1,6-hexanediol promotes Pfk2-GFP aggregation in *snf1Δ* cells subjected to hypoxia, although this mode of aggregation may differ from normal G body formation. Similar results have been reported with extended treatments of high concentrations of 1,6-hexanediol for yeast P-bodies (Wheeler, Matheny, Jain, Abrisch, & Parker, 2016). Together, these findings suggest that G bodies are largely insensitive to 1,6-hexanediol and thus similar in physical properties to yeast stress granules.

Results from experiments with 1,6-hexanediol may not fully explain the physical properties of a granule. If G bodies behave as liquids, they would be expected to fuse as has been demonstrated for a variety of granules such as P bodies (Kroschwald et al., 2015). Consistent with fusion in G body biogenesis, yeast cells display multiple Pfk2-GFP foci at early time points in hypoxia but only a single focus at later timepoints (Jin et al., 2017). To directly test G body fusion *in vivo,* we mated a and α yeast cells, each expressing Pfk2-GFP, and imaged the mating cells over time following hypoxic incubation. When cells were cultured in hypoxia for 18 hours and subsequently placed under a coverslip, we could track individual G bodies over several hours (Fig S7A). G bodies moved throughout the cell (Fig 6A-B, Sup movies 1-2). When a G body from one mating cell type met a G body from the other mating cell, they were able to fuse into one single G body in the mated diploid cell. Fusion could take as little as two minutes or as long as 60 minutes to resolve into a single focus with a median of 18 minutes from initial contact to fusion to form a single G body (Fig 6A). Although some G bodies stayed oblong when fused, others became more spherical over tens of minutes, suggesting that they behaved as gels with fusion over long timescales. In accordance with these results, we also identified some cells that formed G bodies *de novo* during imaging. Initially, two distinct foci appeared from a diffuse cytoplasm, but over 60 minutes, these foci became brighter and fused rapidly within two minutes of contact (Fig 6B).

**Figure 6:**
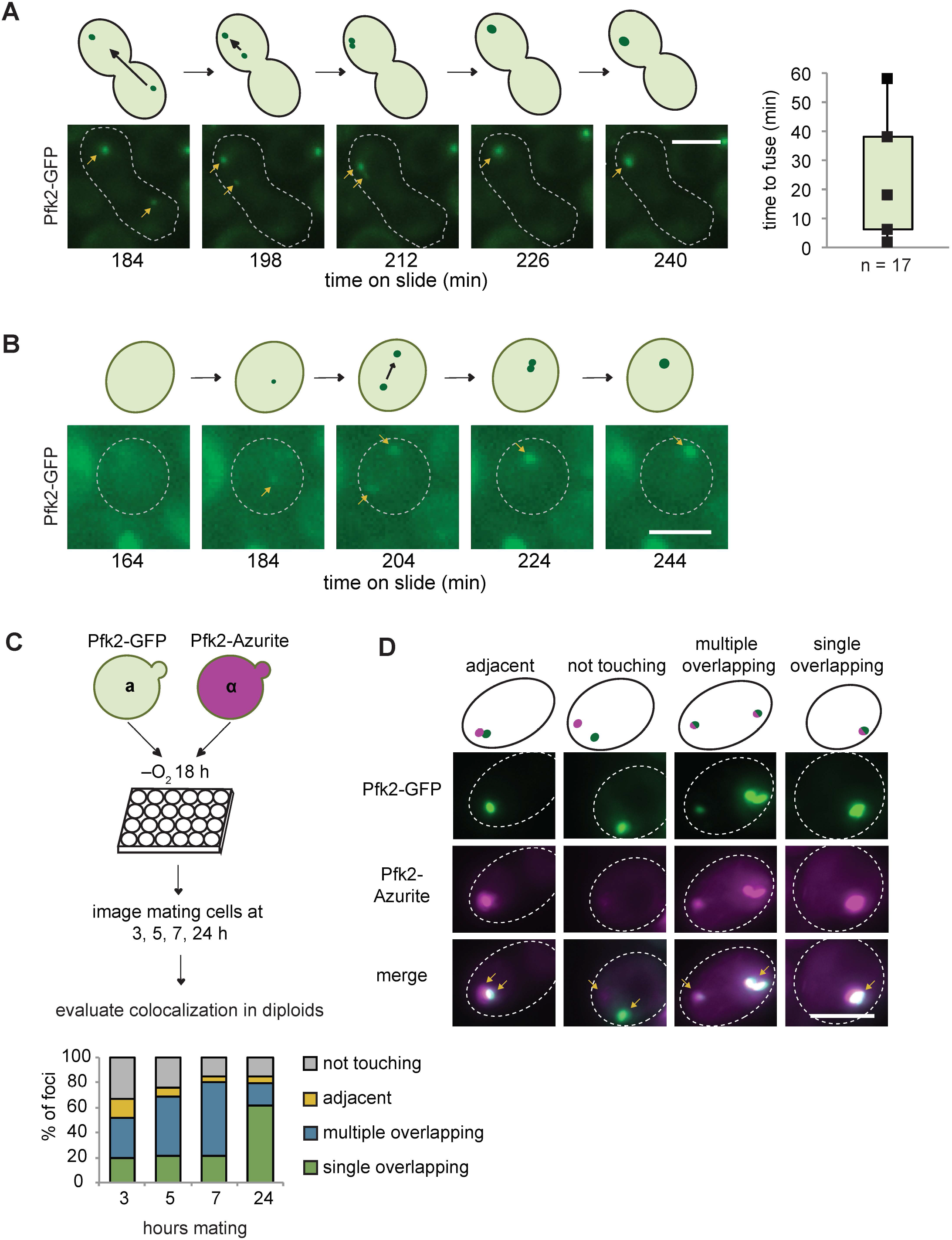
G bodies fuse *in vivo.* (A-B) Still images from time-lapse imaging of Pfk2-GFP in mating cells with cartoon. a and α cells each expressing Pfk2-GFP were cultured together in hypoxia for 18 h, mounted on a slide, and imaged every 2 min for 3–4 h. (A) G-body fusion. Initially, foci are present in opposite ends of mating cell. One G body crosses to the other end and gradually fuses with other G body. (Right) Quantification of time to fuse for multiple G body fusion events from initial contact to fusion. (B) *de novo* G-body formation. Initially, Pfk2-GFP is diffuse in highlighted cell. Two small puncta appear and gradually fuse into one larger focus, becoming more intense with time. (C) Mating for quantification of G body fusion. a and α cells expressing Pfk2-GFP and Pfk2-Azurite, respectively, are grown in hypoxia together and allowed to settle in 24 well plates. Cultures are sampled at 3, 5, 7 and 24 hours and phenotypes are evaluated. (Bottom) Quantification of G body fusion at each time point. Foci were classified as single foci in cells that were overlapping, multiple overlapping foci in cells, adjacent foci in cells, or foci not touching or associated in cells. Data represent mean of three independent experiments (N > 40 foci per timepoint per replicate). (D) (Top) Representative images of mating Pfk2-Azurite and Pfk2-GFP cells displaying different colocalization phenotypes. Arrows indicate G bodies. All scale bars 5 μm.

To gain a better understanding of the frequency of fusion events, we mated a cells expressing Pfk2-GFP to α cells expressing Pfk2-Azurite following hypoxic treatment and allowed them to settle in 24-well plates. Over successive timepoints (3, 5, 7 and 24 h), we measured the fraction of foci with each label that were overlapping, adjacent to, or distinct from puncta with the other label (Fig 6C). These cells would be in a hypoxic environment allowing for some *de novo* granule formation. We observed a high frequency of overlapping puncta, which increased over time. Overlapping puncta likely arose from fusion, although we cannot rule out that a subset arose from *de novo* granule formation in mating cells. The fraction of mating cells with adjacent and unassociated puncta decreased from 41% after 3 h mating to only 11% after 24 h mating, whereas the fraction with overlapping or fused puncta increased (Fig 6D, S7B). The progression from many particles to a single particle is consistent with fusion of granules. Taken together, these results suggest that G bodies can fuse over minutes and are largely insensitive to 1,6-hexanediol, properties reminiscent of gels.

## Discussion

### Physical Properties of G bodies

Here we report that G bodies have properties of phase separated gels *in vivo.* First, similar to yeast and mammalian stress granules, G bodies lack membranes and contain chaperones as well as proteins with intrinsically disordered regions (S. Jain et al., 2016; Jin et al., 2017). Second, we found that G bodies were much less sensitive to 1,6-hexanediol than liquid phase separated structures like P bodies, extending their similarities to stress granules. However, their size appears reduced after 1,6-hexanediol treatment, indicating partial dissolution of the granules. This may be reminiscent of the “dynamic shell” model proposed for stress granules in which a fluid shell surrounds a solid core structure (S. Jain et al., 2016), although there is currently no direct evidence of substructure within G bodies. Third, while G bodies can fuse *in vivo,* a property of liquids (Boeynaems et al., 2018; Elbaum-Garfinkle et al., 2015; Hyman et al., 2014), this fusion takes place on the order of tens of minutes, unlike fusion of P bodies, which takes only seconds (Kroschwald et al., 2015). Similar to P bodies, P granules in the *C. elegans* germline and stress granules can undergo fission and fusion (Brangwynne et al., 2009; Kedersha et al., 2005; Ohshima, Arimoto-Matsuzaki, Tomida, Takekawa, & Ichikawa, 2015). Indeed, formation of G bodies when cells are transitioned to hypoxia involves fusion of smaller granules into larger structures. Early fusion events were insensitive to 1,6-hexanediol due to the presence of single foci even after extended 1,6-hexanediol treatment during hypoxia. Additionally, even after extended periods in hypoxia, G bodies fuse in mating cells, demonstrating that “mature” G bodies can still fuse. Fourth, in contrast to other RNP granules, G bodies are remarkably stable. Like stress granule cores, which can be isolated and are stable in a lysate, G bodies can be purified intact (S. Jain et al., 2016; Jin et al., 2017). However, unlike stress granules, which can disperse on the order of tens of minutes following removal of stress, and nucleoli, which rapidly assemble and disassemble during mitosis (Brangwynne et al., 2011; Feric et al., 2016; S. Jain et al., 2016; Tsai, Ho, & Wei, 2008; Walters, Muhlrad, Garcia, & Parker, 2015), G bodies can persist for tens of hours as revealed in our experiments inducing Pfk2-MqsR-Flag following hypoxic incubation. One possibility is that G bodies initially form from a liquid state and gradually solidify, becoming more similar to protein aggregates of FUS and IDRs of other proteins over time (Lin, Protter, Rosen, & Parker, 2015; Mateju et al., 2017; Murakami et al., 2015). However, G bodies, once formed, can be disrupted by the targeting of an RNase, demonstrating that they are not static. The effect of Pfk2-MqsR-Flag induced after hypoxic treatment (when G bodies have already formed) was largely independent of the concentration of CuSO_4_ added. Thus, a small amount of RNA degradation was sufficient for G body fragmentation. These fragmented foci appeared larger than single puncta in vector control cells, suggesting additional Pfk2-GFP could accumulate in these puncta in the absence of RNA. This is consistent with RNA primarily being required for G body fusion and nucleation. It is unclear whether resident G-body proteins ever return to the soluble cytosolic pool when cells are shifted back from hypoxic to normoxic conditions.

Similar to proteins in other granules, protein recruitment to G bodies relies on multivalent interactions. Both the N- and C-terminal domains of Pfk2 are necessary but not sufficient for G-body recruitment. Pfk1 is recruited to G bodies via an interaction with Pfk2. Additionally, protein-RNA interactions are required for G body formation. Such multivalent interactions are necessary for phase separation and can drive granule formation (Hyman et al., 2014; Li et al., 2012).

### RNA in phase separation

By targeting RNase fusion proteins to sites of G body formation, like a Trojan Horse we show that RNA is required for G body formation *in vivo*. Degradation of RNA in existing G bodies led to multiple puncta, suggesting that RNA is required for G body stability or integrity. Our mass spectrometry analysis of the “RBPome” identified hundreds of mRBPs, including glycolysis enzymes, consistent with previous work (Matia-González et al., 2015) (Beckmann et al., 2015.; Scherrer, Mittal, Janga, & Gerber, 2010). By PAR-CLIP-seq, a comprehensive picture of an extensive glycolysis enzyme-bound transcriptome is emerging. Intriguingly, these enzymes bind similar transcripts primarily in the 3’ UTRs and coding regions of their substrate mRNAs. We identified a common set of transcripts bound by multiple glycolysis enzymes in normoxic conditions. We determined, by RIP-qPCR, that at least some of these mRNA substrates were then recruited along with their binding protein partners to G bodies during formation. Taken together with the requirement for RNA at the site of G body formation, we propose a model in which RNA serves as a scaffold for G body nucleation and growth. Consistent with this model, addition RNA promotes the aggregation of the glycolysis enzyme Cdc19 *in vitro* (Saad et al., 2017). Among the commonly bound transcripts were a number of mRNAs encoding the glycolysis enzymes themselves. Weak, dynamic interactions between G body components with RNA and with each other would allow for the growth of G bodies. It has been proposed that RNA interactions with glycolysis enzymes can facilitate post-transcriptional regulation of the pathway. This mode of regulation could additionally contribute to G body formation through concentration of nascent proteins due to spatially segregated translation of glycolysis enzyme mRNAs. Interactions with glycolysis enzyme mRNAs may specifically facilitate multivalent interactions of glycolysis enzymes with RNA by providing a common set of substrates.

### Phase separation in the control of metabolic pathways

Although spatial organization of pathways to concentrate constituent enzymes is not novel, phase separation is emerging as a new mechanism to achieve this type of organization. Even for glycolysis, pathway compartmentalization is known to occur; glycolysis enzymes are concentrated in membrane-bound compartments called glycosomes in various protozoa, including in *Trypanosoma brucei,* which can survive in anaerobic conditions (Michels, Bringaud, Herman, & Hannaert, 2006; Opperdoes, 1987). However, phase separation is mechanistically distinct in that it does not require formation of a membrane or specific transporters. G bodies represent an addition to the known metabolic pathways organized by phase separation mechanisms. Three recent studies have demonstrated glycolysis enzyme coalescence in hypoxia, which is associated with increased rates of glucose flux (Jang et al., 2016; Jin et al., 2017; Miura et al., 2013). Thus, G bodies and glycosomes may represent a case of convergent evolution. Similar structures also form in the neurons of *C. elegans,* where enzyme clustering in response to hypoxia was associated with proper synaptic function, suggesting that glycolysis enzymes coalesce to increase glycolysis and meet the local high energy demand during synapses (Jang et al., 2016). Furthermore, cancer cell lines form small aggregates of glycolysis enzymes even in the presence of oxygen, suggesting that concentrating glycolysis enzymes is a highly conserved process (Kohnhorst et al., 2017). Additionally, pyrenoids, which enhance the rate of carbon fixation in *Chlamydomonas* by concentrating CO_2_ in the presence of ribulose 1,5 bisphosphate carboxylase oxygenase (RuBisCo), were recently shown to form via phase separation (Freeman Rosenzweig et al., 2017). Carbon metabolism, then, can be controlled both at the level of fixation and harvesting via phase separation into liquid compartments.

The precise mechanism of enhanced glycolysis activity is unclear in G bodies. Recent *in vitro* measurements of dextranase activity in artificial liquid-liquid phase separated compartments suggest that phase separation can enhance reaction rates through decreased substrate inhibition (Kojima & Takayama, 2018). The G body resident factor phosphofructokinase mediates one of the irreversible steps in glycolysis. It is also subject to substrate inhibition by ATP and may experience increased specific activity akin to release from substrate inhibition when dextranase undergoes liquid-liquid phase separation. However, purinosomes, which are complexes composed of purine biosynthesis enzymes, are thought to enhance pathway activity through substrate channeling (An, Kumar, Sheets, & Benkovic, 2008; Zhao, French, Fang, & Benkovic, 2013). Concentrating enzymes in RNP granules may achieve similar results or enhance reaction flux rates via concentrating intermediate metabolites. Alternatively, concentration of energy producing enzymes with their cognate mRNAs may facilitate enhanced translation of the glycolysis enzymes, thus promoting pathway activity without increasing specific activity of the enzymes. Understanding the mechanism and function of metabolic pathway enhancement by phase separation will require future mechanistic studies.

## Methods

### Yeast strains and culturing

Strains used are listed in Supplemental Table 3. Deletion mutants and integrated transgenic strains were generated through transformation of amplified auxotrophic markers at the site of PCR amplified fragments with at least 40 nt of overlapping sequence to 3’ and 5’ UTR sequences. GFP mutants were derived from the yeast GFP tagged library (Huh et al., 2003). TAP tagged mutant strains were derived from the yeast TAP library (Ghaemmaghami et al., 2003). Plasmids were transformed with standard lithium acetate transformation (Gietz & Woods, 2002) and selected on SD with appropriate amino acid supplements and auxotrophic selection.

Cells were grown as indicated either in YPD (2% peptone, 1% yeast extract, 5% glucose) or SMD (0.67% yeast nitrogen base, 5% glucose, appropriate amino acid and vitamin supplements). For hypoxic growth of yeast for imaging, cells were reinoculated from stationary phase starter cultures in indicated media in 0.5-1 ml volumes 24-well plates at an OD of 0.05. Cells were grown for indicated amounts of time in hypoxia using the AnaeroPack system in 2.5 L boxes (Mitsubishi Gas Chemical). For larger-scale biochemical experiments, cells were grown in 125 ml Erlenmeyer flasks in 10–25 ml of media in 7 L boxes. For analysis of copper inducible proteins, cells were grown in SMD lacking uracil and supplemented with CuSO_4_ at varying concentrations.

### Glucose starvation and 1,6-hexanediol treatment

For glucose starvation, cells were reinoculated from YPD starter cultures into SMD at an OD of 0.15. Cells were grown for 6 h shaking at 30° C to log phase and pelleted for 2 min at 500 X G. Cells were washed and resuspended in SM media lacking glucose with or without 5% 1,6-hexanediol for 30 min at room temperature. Cells were either immediately imaged or had 1,6-hexanediol in PBST directly added to the media, were incubated for 1 h at room temperature and then imaged.

For hypoxic treatment with 1,6-hexanediol, cells were grown in YPD and reinoculated in SMD with 0%, 1%, or 2% 1,6-hexanediol at an OD of 0.05 and grown 20 h in hypoxia and imaged. For treatment with 1,6-hexanediol after hypoxia, cells were cultured in the same way and had 1,6-hexanediol in PBST directly added to the media, incubated for 1 h at room temperature, and then imaged.

### Plasmid construction

Plasmids were generated by overlap extension PCR fusing fragments amplified from genomic DNA to fragments amplified from plasmid sources. Pfk2-Rnase A was generated by fusing the Pfk2-linker sequence form THY62 to RNase A, omitting the signal peptide from pet22B RNase A. pET22b RNase A was a gift from Ronald Raines (Addgene plasmid # 58903). Pfk2-MqsR was generated by fusing the Pfk2-linker sequence from THY62 to MqsR amplified with a 3’ 1X Flag peptide sequence from pSLC-241. pSLC-241 was a gift from Swaine Chen (Addgene plasmid # 73194). Pfk2-MqsR-MqsA was generated by fusing Pfk2-MqsR-Fl to MqsA and subsequently fusing the Pfk2 3’ UTR. Cytoplasmic MqsR was generated by amplifying MqsR from pSLC-241. Each construct was subsequently fused to the Pfk2 3’ UTR amplified from genomic DNA and subcloned into pCu416CUP1 (ATCC^®^ 87729^TM^) using Spe1 and Xho1 sites. Pfk2-Rnase A^H12A^ was introduced with site-directed mutagenesis and verified by sequencing.

### Yeast immunofluorescence

Yeast immunofluorescence protocol was adapted from (Amberg, Burke, & Strathern, 2005). Briefly, cells were grown in hypoxia for indicated periods of time to an OD of ~2 in 1 ml volumes. Formaldehyde was added to culture media to 1% for 10 min and then washed away. Cells were resuspended in KM (50 mM potassium phosphate, 5 mM MgCl_2_, pH 6.5) with 4% formaldehyde for 1 h at 30° C and washed twice with KM and once with KM with 50% sorbitol (KMS). Cells were resuspended in 50 μL KMS with 5 U Zymolyase (Zymo E1004) for 20 min at 37 °C. Cells were washed and resuspended in KMS and adhered to Superfrost Plus™ (Fisher 12-550-15) slides. For probing Pfk2-Rnase A, cells were dipped in ice-cold methanol for 6 minutes followed by acetone for 30 seconds and dried at 50 °C. For Pfk2-MqsR, to preserve GFP fluorescence, this step was omitted. Cells were blocked in 1% BSA in PBST 2 h at room temperature. For Pfk2-Rnase A, cells were incubated with 1:200 rabbit anti-S tag (Genscript A00625) overnight and 1:1000 mouse anti GFP (Thermo Fisher A-11120) for Pfk2-Rnase A and 1:200 mouse anti Flag in PBST overnight at 4 °C. Cells were washed three times in PBST and 1:500 Donkey anti-mouse Alexa 488 Alexa Fluor (A-21202), and 1:500 Goat anti Rabbit Alexa Fluor 647 (A-21245) for Pfk2-Rnase A in PBST was added for 1 h at room temperature. For Pfk2-MqsR-Fl experiments, 1:500 Donkey anti-Mouse Alexa Fluor 647 (A-31571) was added in PBST for 1 h at room temperature. Cells were mounted in Vectashield plus DAPI (Vector laboratories, H1200) and imaged.

### Yeast fluorescence microscopy imaging and analysis

All cells were imaged using a Zeiss AxioImager M2 with an ORCA-Flash 4.0 LT camera with band pass GFP filter (Zeiss 38 HE eGFP) illuminated with a 488 nm LED with either 40X or 100X objective taking Z stacks to cover the entire cell. Immunofluorescence images were taken with illumination from a mercury halide arc lamp and band pass GFP filter and CY5 filter (Zeiss 50).

For assaying G-body formation, cells were manually counted and classified into one of three categories: cells with single puncta, cells with multiple puncta, and cells with no puncta. At least 100 cells were considered for each replicate and condition.

For mating experiments, cells of each mating type were mixed at an OD of 0.05 and grown 18 h –O_2_. Cells were allowed to settle in 24-well plates for mating to proceed and sampled after 3, 5, 7, and, 24 h. For mating cells with Pfk2-Azurite and Pfk2-GFP, Azurite was imaged with a BFP band pass filterset (Zeiss 96 HE) illuminated with a 365 nm LED, and GFP was imaged as above. To compensate for high background, mating cells were resuspended in PBS before imaging. Z stacks were taken in each channel and brightfield. Overlapping puncta were manually counted in all Z planes, and puncta were classed into four categories: overlapping puncta in cells with one focus, overlapping puncta in cells with multiple foci, adjacent puncta, and puncta not associated with other puncta. For mating cells with Pfk2-GFP only to observe kinetics, cells were mixed and grown 18 h –O_2_ and placed on a slide. Fields of cells were imaged taking Z stacks every 2 minutes through a GFP filter.

For size distributions of G bodies, local maxima were identified in maximum intensity projections in FIJI (Schindelin et al., 2012) using the Find Maxima tool. A square with sides of 4.9 μm was drawn around each maximum. Average background from 3 separate spots in each image was subtracted. Using custom Python scripts, each focus was fit using nonlinear least squares to a 2-dimensional Gaussian distribution of the form:

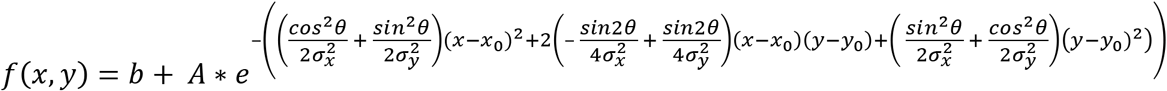

where b is the baseline fluorescence, A is the amplitude of the peak fluorescence of the focus, x_0_ and y_0_ define the center coordinate, σ_x_ and σ_y_ represent the standard deviation along each axis and θ defines the rotation of the punctum.

### Yeast PAR-CL-Mass Spectrometry

3 L of BY4742 were cultured from an initial OD600 of 0.003 and grown until 0.7-0.8. Cycloheximide was added to a final concentration of 0.1 mg/ml and incubated at 30°C for 5 min. Cells were pelleted by centrifugation for 5 min at 4,000 rpm at 18°C using a JLA-10.5 rotor in an Avanti J-26XP centrifuge, resuspended in 10 ml 1XPBS (with 0.1 mg/mL cycloheximide), transferred to a 150-mm glass Petri dish, placed on ice, and irradiated 4 times with 365-nm UV light at 150 mJ/cm^2^ using a UVP CL-1000L UV crosslinker. The cells were then transferred to a 15 mL conical tube and pelleted for 3 min at 3,000Xg at room temperature. After removing the PBS, the cells were frozen in liquid nitrogen. For the negative control, cells were frozen without UV irradiation. Frozen cell pellets were pulverized for two cycles, each for 1 min, at 30 Hz on a Retsch MM 4000 ball mill homogenizer. Sample chambers were pre-chilled in liquid nitrogen and re-chilled between cycles. The resulting frozen powdered homogenate was resuspended in 3 mL of polysome lysis buffer (20 mM HEPES, pH 7.5, 140 mM KCl, 1.5 mM MgCl2, 1% Triton X-100, 1X cOmplete Mini Protease Inhibitor, EDTA free, 0.1 mg/mL cycloheximide) and incubated on ice for 10 min. Cell debris was removed by centrifugation for 2 min at 3,000Xg at 4°C. The supernatant fraction was clarified by a 20,000Xg spin for 10 min at 4°C and supplemented with 12 μL SUPERase·In (20 U/μL). 0.3 mL of 1 M (34.2% w/v) sucrose cushion solution, prepared in polysome lysis buffer, was layered on the bottom of each 11 × 34-mm polycarbonate centrifugation tube. 3 mL of clarified lysate was loaded onto 3 sucrose cushions (1 mL per cushion) and spun for 80 min at 54,000 rpm at 4°C in a TLS-55 rotor using an Optima Ultracentrifuge MAX-E. After centrifugation, the top 1 mL of solution was recovered from each tube (3 ml in total), mixed with 1.5 g of guanidine thiocyanate (GuSCN, Promega V2791), vortexed to dissolve GuSCN, and heated for 5 min at 65°C. A Zeba desalting column (7K MWCO, 10 mL, Pierce 89894) was used to remove GuSCN and to exchange buffer to 50 mM NaCl buffer (20 mM HEPES, pH 7.3, 50 mM NaCl, 0.5% Sarkosyl, 1 mM EDTA). Buffer-exchanged lysate was combined with 0.1 vol of 5 M NaCl to adjust the salt concentration to 0.5 M and supplemented with 6 μL SUPERase·In (20 U/μL). The lysate was incubated with 37.5 mg of oligo(dT)25 beads (NEB S1419S) for 30 min at 4°C on a Nutator. Beads were washed four times with ice-cold low-salt wash buffer (20 mM HEPES, pH 7.3, 0.2 M NaCl, 0.2% Sarkosyl, 1 mM EDTA). RNAs were eluted in 1 mL of elution buffer (10 mM HEPES, pH 7.3, 1 mM EDTA) by heating for 3 min at 65°C. Eluted RNAs were concentrated to 80 μL with Amicon spin filters (3 KD cutoff, Millipore UFC500324). Concentrated RNAs were mixed with 40 μL of 3X SDS sample buffer (150 mM Tris, pH 6.8, 6% SDS, 30% glycerol, 3% beta-mercaptoethanol, 37.5 mM EDTA, 0.06% Bromophenol blue) and heated for 5 min at 65°C. 20 μL of sample was loaded per lane (6 lanes total) of a 4-12% NU-PAGE Bis-Tris gel and run for 10 min at 100 V, followed by 70 min at 150 V. The gel was stained with Colloidal Blue (Invitrogen LC6025), and a gel piece, 0.1 cm-1.0 cm below the well, was excised and stored at −80°C before mass spectrometry analysis.

For mass spectrometry, unless otherwise noted, all chemicals were purchased from Thermo Fisher Scientific (Waltham, MA). Deionized water (18.2 MW, Barnstead, Dubuque, IA) was used for all preparations. Buffer A consists of 5% acetonitrile, 0.1% formic acid; buffer B consists of 80% acetonitrile, 0.1% formic acid; and buffer C consists of 500 mM ammonium acetate. All buffers were filtered through 0.2-mm membrane filters (PN4454, Pall Life Sciences, Port Washington, NY). In-gel digestion was performed as in (Jensen, Wilm, Shevchenko, & Mann, 1999) with the following adjustments: Gel particles were rehydrated with 10 mM Tris (2-carboxyethyl) phosphene in 100 mM NH4HCO3 and incubated for 30 min at room temperature (RT). Digestion buffer was 50 mM NH4CO3, 5 mM CaCl2, containing 12.5 ng/μL trypsin. The gel pieces were rehydrated at RT for 30–45 min. The enzyme supernatant fraction was not removed, and 50 μL digestion buffer, without enzyme, was added before overnight digestion. A MudPIT microcolumn (Wolters, Washburn, & Yates, 2001) was prepared by first creating a Kasil frit at one end of an undeactivated 250-mm ID/360-mm OD capillary (Agilent Technologies, Inc., Santa Clara, CA). The Kasil frit was prepared by briefly dipping a 20-to 30-cm capillary in well-mixed 300 mL Kasil 1624 (PQ Corporation, Malvern, PA) and 100 mL formamide, curing at 100°C for 4 h and cutting the frit to ~2 mm in length. Strong cation exchange particles (SCX Partisphere, 5 mm dia., 125 Å pores, Whatman) were packed inhouse from particle slurries in methanol to 2.5 cm. 2.5-cm reverse phase particles (C18 Aqua, 3 mm dia., 125 Å pores, Phenomenex, Torrance, CA) and were then packed into the capillary using the same method as SCX loading, to create a biphasic column. The MudPIT microcolumn was equilibrated using 60% buffer A, 40% buffer B for 5 min, and followed by 100% buffer A for 15 min. An analytical RPLC column was generated by pulling a 100 mm ID/360 mm OD capillary (Polymicro Technologies, Inc, Phoenix, AZ) to 5 mm ID tip. Reverse phase particles (Aqua C18, 3 mm dia., 125 Å pores, Phenomenex, Torrance, CA) were packed directly into the pulled column at 800 psi until they were 12-cm long. The column was further packed, washed, and equilibrated with buffer B followed by buffer A. The MudPIT microcolumn was connected to an analytical column using a zero-dead volume union (Upchurch Scientific (IDEX Health & Science), P-720-01, Oak Harbor, WA). LC-MS/MS analysis was performed using an Eksigent nano-flow pump and a Thermo LTQ-Orbitrap using an in-house-built electrospray stage. MudPIT experiments were performed where each step corresponds to 0, 10, 20, 30, 40, 50, 60, 70, 80, 90, and 100% buffer, C being run for 5 min at the beginning of each gradient of buffer B. Electrospray was performed directly from the analytical column by applying the ESI voltage at a tee (150 mm ID, Upchurch Scientific) while flowing at 350 nL/min through the columns. Electrospray directly from the LC column was done at 2.5 kV with an inlet capillary temperature of 250°C. Data-dependent acquisition of MS/MS spectra with the LTQ-Orbitrap were performed with the following settings: MS/MS on the 10 most intense ions per precursor scan, 1 microscan, unassigned and charge state 1 reject; dynamic exclusion repeat count, 1, repeat duration,-30 second; exclusion list size 120; and exclusion duration, 120 second. Tandem mass spectra were extracted from raw files using RawExtract 1.9.9 (McDonald et al., 2004) and were searched against a yeast protein database (http://www.yeastgenome.org) with reversed sequences using ProLuCID (Peng, Elias, Thoreen, Licklider, & Gygi, 2003; Xu et al., 2015). The search space included all fully and half-tryptic peptide candidates. Carbamidomethylation (+57.02146) of cysteine was considered a static modification. Peptide candidates were filtered using DTASelect (v2), with the following parameters: –p 1 – y 1 –trypstat –fp 0.01 –extra –DM 10 –DB –dm <in (McDonald et al., 2004; Tabb, McDonald, & Yates, 2002).

### Yeast PAR-CLIP western blotting and autoradiography

RBP validation was performed similar to the PAR-CLIP protocol but omitting the linker ligation steps. Cells collected from 50 mL of log-phase culture were used in each IP. After autoradiography, the same membrane was blotted with Peroxidase Anti-Peroxidase (Sigma-Aldrich P2416) at 1:10,000 dilution and developed with ECL substrates (Pierce 32209).

### Yeast PAR-CLIP-seq (Pfk2, Eno1, Fba1)

PAR-CLIP was performed as described previously (Freeberg et al., 2013). Briefly, yeast were grown to mid-log phase and irradiated with 365-nm UV. Cross-linked cells were lysed, treated with Rnase T1, and mixed with IgG magnetic beads to affinity isolate each TAP-tagged protein. Lysates were then subjected to Rnase T1 digestion, CIP treatment, 3’ DNA linker ligation, 5’ end phosphorylation, and SDS-PAGE. After nitrocellulose transfer, cross-linked RNAs were visualized by autoradiography. Bands corresponding to each protein were excised and incubated with proteinase K. RNAs were collected by centrifugation and loaded onto a 6% TBE UREA gel. Gel pieces corresponding to 70-90 nt RNA were excised followed by amplification of the RNA fragments by RT-PCR. Amplicons were purified, run on a 10% TBE gel, and gel pieces corresponding to 96-116 bp DNA were excised. DNA fragments were amplified by PCR for two rounds and sequenced on an Illumina HiSeq 2000 sequencer. All primers used are as listed previously (Freeberg et al., 2013). Specifically, Indexes 1, 3, and 1 (for Pfk2, Fba1, and Eno1, respectively) barcoded 3’ DNA linker oligos and reverse transcription primers were used.

Sequenced PAR-CLIP-seq read data were processed as described for Puf3 PAR-CLIP-seq (Freeberg et al., 2013). Briefly, reads were processed to remove linkers and sorted into libraries based on six-nucleotide barcodes. Next, reads were removed if they met any of the following criteria: <18 nucleotides, only homopolymer As, missing 3’ adapter, 5’-3’ adapter ligation products, 5’-5’ adapter ligation products, and low quality (more than 4 bases with quality scores below 10 or more than 6 bases with a quality score below 13). High-quality reads were mapped to the S. *cerevisiae* genome (S288C, sacCer3) with Bowtie (Langmead, 2010) using the following parameters: -v 3 (map with up to 3 mismatches), -k 275 (map at up to 275 loci), --best, and –strata.

### Pfk2, Fba1, and Eno1 binding site generation

Reads were assembled into binding sites by aggregating overlapping reads harboring 0–2 T-to-C conversion events. Only binding sites containing at least 1 T-to-C conversion event were considered high-confidence binding sites. For each library, the counts of sequencing reads covering each position within a binding site were averaged and normalized to the total number of millions of mapped reads in that library. To filter off low-coverage binding sites, a reads-per-million (RPM) mapped reads threshold for each library was empirically determined by simulating replicate data from each PAR-CLIP-seq dataset. Two sets of binding site RPM values were randomly sampled from all binding sites passing a minimum RPM threshold in a single dataset such that each sample contained 20% of the binding sites. A non-parametric two-sample Kolmogorov-Smirnov (K-S) test was performed on the two sets of RPM values, and the resulting K-S test statistic was recorded. This test was repeated 10,000 times for each of 36 RPM threshold values ranging from 0 to 25. Mean K-S test statistic values were plotted for each RPM threshold value, and a final binding site RPM threshold value for the library was chosen when the K-S test statistic stabilized (Fig. S1B). For Pfk2p, an empirical RPM threshold of 5 RPM was used, and a threshold of 0.5 RPM was used for Eno1p and Fba1p. After filtering, binding site RPM values were normalized to gene expression RPKM values from previously published data (Freeberg et al., 2013). Binding sites were annotated using custom scripts to known genomic elements in the S288C (sacCer3) yeast genome. ORFs with unannotated UTRs were hierarchically assigned UTRs from the following: (Nagalakshmi et al., 2008; Yassour et al., 2009). GO term analysis was performed using the g:Profiler web server (Reimand et al., 2016).

### G body Immunoprecipitation and RNA extraction

Protein G Dynabeads (ThermoFisher Scientific, 1004D) were conjugated to mouse anti-Flag M2 antibody (Sigma Aldrich F1804) as previously described (Jin et al., 2017). Beads were stored in 10% BSA in lysis buffer (150 mM NaCL, 25 mM Tris-HCl pH 7.5, 5 mM EDTA, 0.5 mM Dithiothreitol, 0.5% NP-40, 1 cOmplete^TM^ mini EDTA-free protease inhibitor cocktail tab per 5 ml (Roche, 0463159001), 40 U RNaseOUT^TM^ (Thermo Fisher 10777019) per ml) until use. Pfk2-GFP-Flag cells BY4742 cells were reinocculated from overnight YPD starter cultures into 125 ml of YPD at an OD of 0.005 split into five 125 ml Erlenmeyer flasks. Cells were grown in hypoxia for 18 h. Cells were imaged to ensure normal G body formation. 100 Ods of cells were spun down at 3,000 X G for 10 min and decanted with remaining media being aspirated. Cells were resuspended in 1.5 ml of lysis buffer and lysed by glass bead lysis for 10 minutes alternating vortexing and ice every 30 seconds. Large cell debris were removed from the supernatant by centrifugation at 500Xg for 5 min. The supernatant was precleared once with 25 μL of Protein G Dynabeads nutating for 30 minutes at 4° C. G bodies were then pelleted by centrifugation at 5,000Xg for 10 min and resuspended in 1 ml of lysis buffer and incubated with Dynabeads for 2 h at 4° C. After 1 hour, Dynabeads were imaged to ensure capture of G bodies. 250 μL of the flow through was saved for RNA extraction. Beads were washed 3X with 1 ml of lysis buffer and once with 1 ml of Proteinase K buffer (100 mM Tris pH 7.5, 50 m NaCL, 10 mM EDTA). Beads were resuspended in 120 μL 2 mg/ml 3X Flag Peptide (Millipore F4799) in Proteinase K buffer and eluted for 30 minutes at 37° C on a thermomixer at 850 RPM. After 15 min, 1 μL of the eluate was imaged to validate elution of G bodies from Dynabeads. The eluate was mixed 1:1 with 8 mg/ml Proteinase K (Thermo Fisher 25530015) in proteinase K buffer for 1 h in a thermomixer at 37° C at 850 RPM. Eluates and flow-through RNA was extracted with Tri Reagent (Sigma-Aldrich T9424) according to the manufacturer instructions. The aqueous phase was mixed 1:1 with isopropanol with 1 μL of glycogen (Thermo Fisher R0561) and precipitated for 4 hours at – 80° C. RNA was pelleted by centrifugation at 4° C at 21,000Xg for 30 minutes. RNA was washed 3 times with 80% ethanol and re-precipitated in 80% ethanol with sodium acetate overnight at −80° C. RNA was pelleted again and washed again and dissolved in 30 μL of H_2_O. For each sample, 10 μL of RNA was used to generate two technical replicates of cDNA with the High-Capacity cDNA reverse transcription Kit (Thermo Fisher 4368813) per the manufacturer’s instructions. Individual probes (Table S2) were used to measure different RNA species using Absolute Blue qPCR SYBR Green (Thermo Fisher AB4166B) per the manufacturer instructions with a Bio-RAD CF96 Real Time PCR thermal cycler. As the flow through RNA represented 25% of the total flow through, 2 cycles were subtracted from the resulting Cq values. For each probe, the ΔCq of elution RNA – flow through RNA was measured and the percent of input was plotted.

## Acknowledgements

We thank all members of the Kim Lab (Amelia Alessi, Charlotte Choi, Mindy Clark, Jessica Kirshner, Nathan Roach, Alex Rittenhouse, and Rebecca Tay) for helpful suggestions. Additionally, we thank Sua Myong and Himani Galagali for comments on the manuscript. This work was supported by a grant from the NIH (RO1GM129301).

---

**Figure S1:**
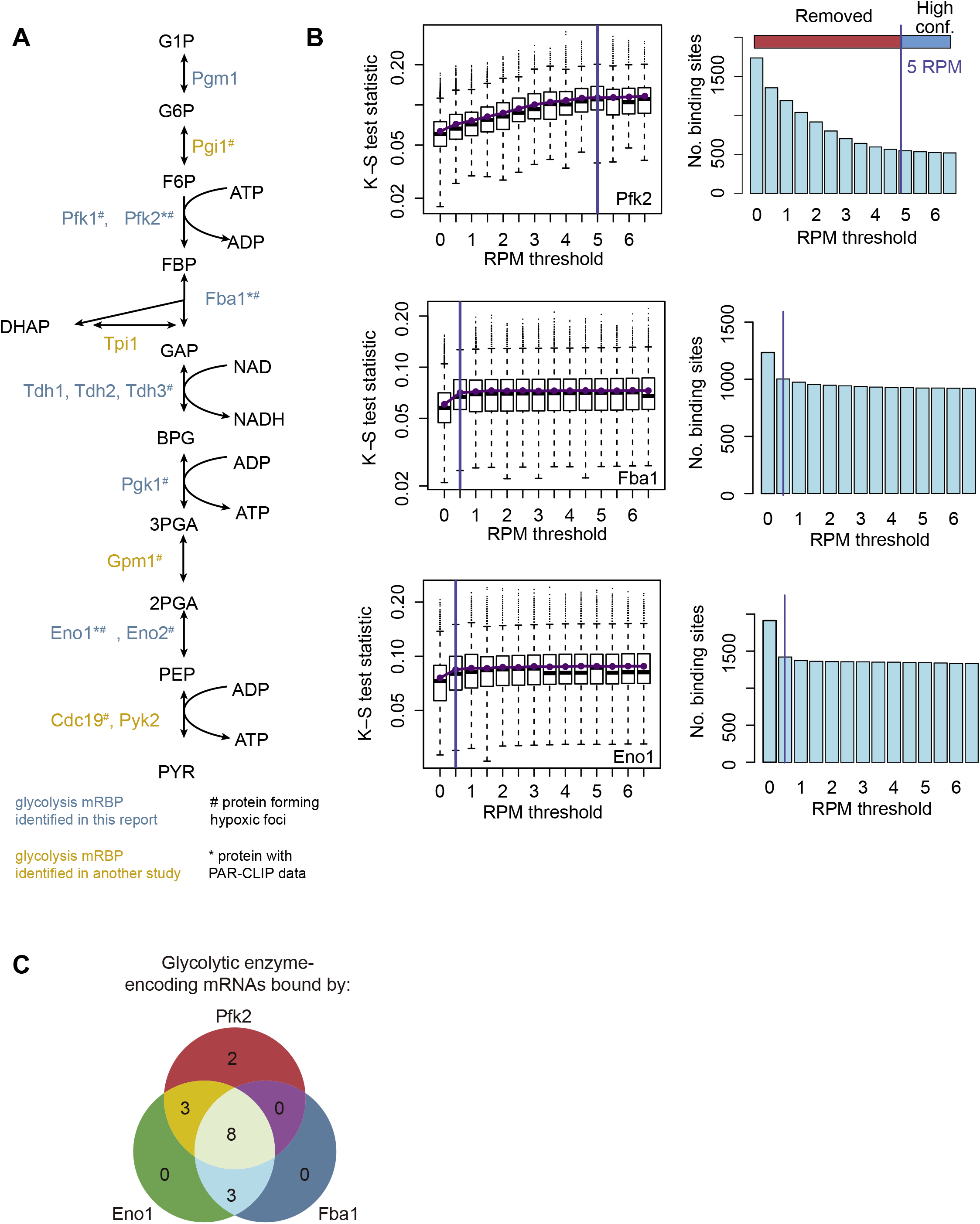
Related to Figure 1. Identification of Glycolysis enzyme binding sites. (A) Glycolytic enzymes indicated in blue were identified in the dataset in this report among yeast mRBPs. Glyclolytic enzymes indicated in yellow were identified in (Beckmann et al., 2015) and/or (Castello et al., 2012). # indicates glycolysis enzymes with observed punctate localization in hypoxia (Jin et al., 2017; Miura et al., 2013). * Indicates PAR-CLIP analysis of protein appears in this report. (B) Empirical determination of high-confidence RPM thresholds of 5 RPM for Pfk2 binding sites from PAR-CLIP-seq. Thresholds of 0.5 RPM were determined for sites bound by Eno1 and Fba1. (C) Sixteen glycolytic enzyme-encoding mRNAs are bound by Pfk2, Eno1, and/or Fba1.

**Figure S2:**
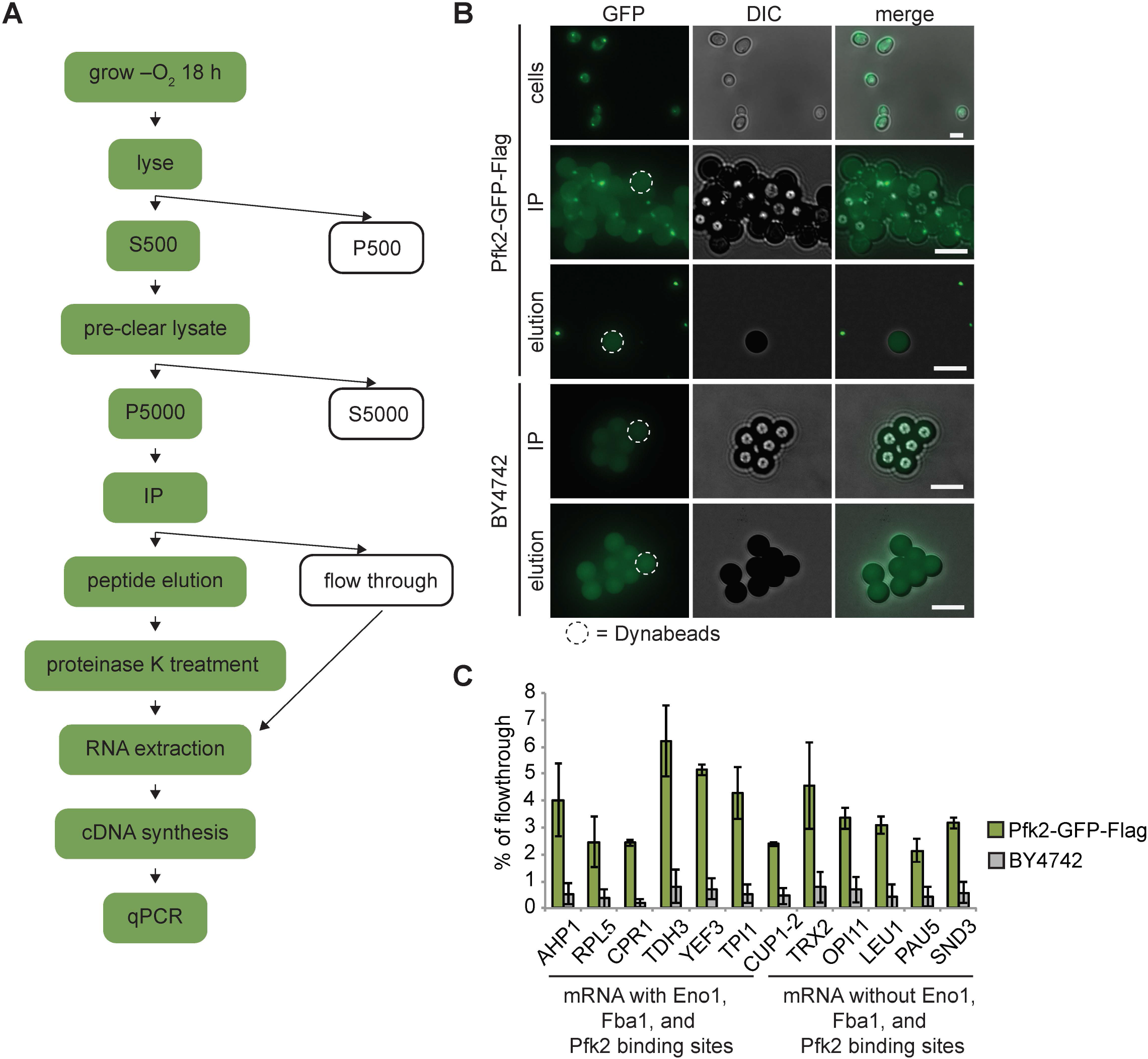
Related to Figure 2. G bodies co-purify with RNA. (A). Schematic of G body RIP purification. Green colored steps indicate presence of G bodies. (B). Images from various steps of purification. Cells expressing Pfk2-GFP-Flag form G bodies. In immunoprecipitations with Pfk2-GFP-Flag, large foci (small brightly fluorescent structures) are present on the surface of Dynabeads (large weakly fluorescent structures) while there are no foci in immunoprecipitations of BY4742 lysates. Elution with flag peptide releases foci from the surface of Dynabeads allowing recovery of G bodies. Dotted outline represents an example of a Dynabead in each image with Dynabeads. All scale bars are 5 μm (C). qPCR of 12 mRNAs. Plotted is the percent of RNA in eluted with G bodies of the flowthrough recovered in RIP samples for 3 biological replicates. AHP1, RPL5, CPR1, TDH3, YEF3 and TPI1 mRNAs have binding sites for glycolysis enzymes identified by PAR-CLIP. CUP1-2, TRX2, OPI11, LEU1, PAU5 and SND3 do not.

**Figure S3:**
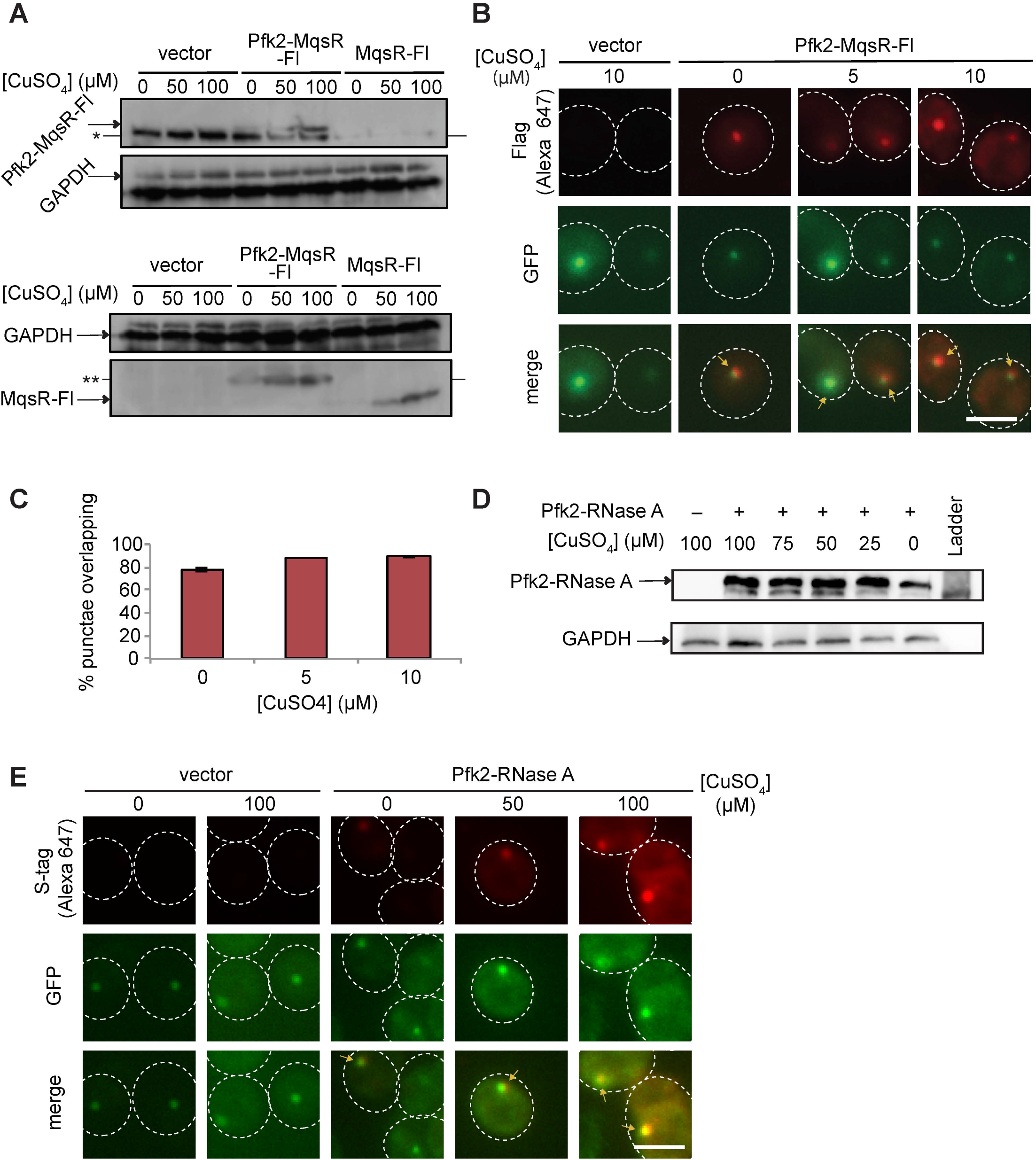
Related to Figure 3. Validation of RNase tagged Pfk2 variants. (A). Western blot showing induction and expression of Pfk2-MqsR-Fl and MqsR-Fl. Top blot was from an 8% polyacrylamide gel. Bottom blot was from a 12% polyacrylamide gel. GAPDH served as a loading control. Nonspecific band of approximately 100 kDa is present in all samples, whereas Pfk2-MqsR-Fl is present at 110 kDa. MqsR-Fl is about 12 kDa. MqsR-Fl and Pfk2-MqsR-Fl were probed with a monoclonal anti-flag antibody. * indicates nonspecific band. ** indicates degradation product. (B) Immunofluorescence of Pfk2-MqsR-Fl (red) probed by a monoclonal anti-flag antibody shows colocalization with endogenous Pfk2-GFP signal (green). No Flag signal is present in vector control. (C) Quantification of Flag foci overlapping Pfk2-GFP foci. Data represent mean and standard deviation of two replicates (N > 100 per replicate). (D) Western blot of cells expressing Pfk2-RNase A at different concentrations of CuSO_4_ showing induction. Pfk2-RNase A was detected by a rabbit anti-S-tag antibody. GAPDH served as a loading control. (E) Immunofluorescence showing colocalization of Pfk2-RNase A (red) with Pfk2-GFP (green). GFP was probed with a monoclonal anti-GFP antibody. RNase A was probed with a rabbit anti-S-tag antibody. Arrows indicate G bodies. All scale bars are 5 μm.

**Figure S4:**
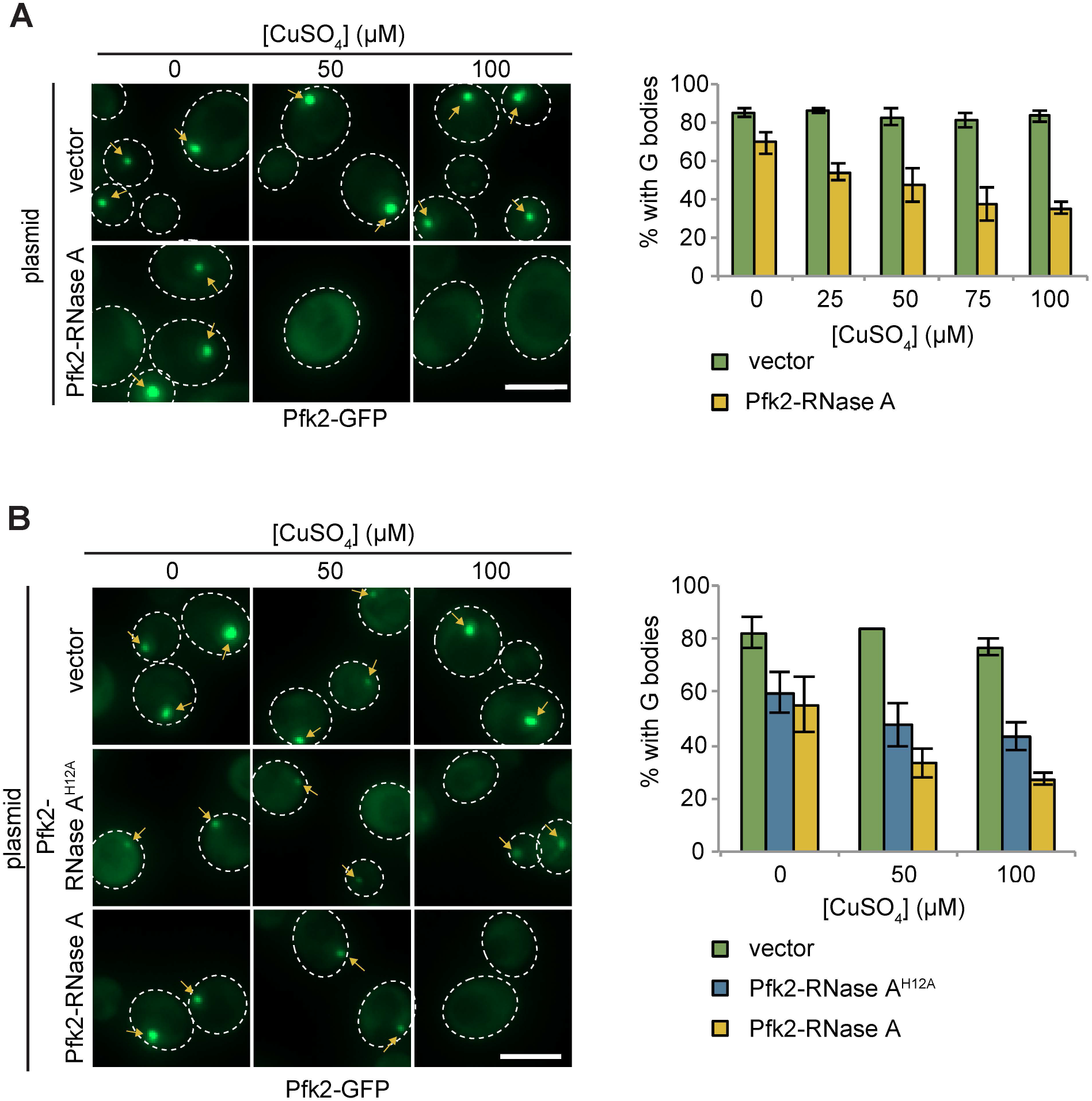
Related to Figure 3. Pfk2-RNase A inhibits G-body formation. (A). (Left) Representative images of hypoxic Pfk2-GFP localization with varying concentrations of CuSO_4_ in cells with a vector control or plasmid expressing Pfk2-RNase A lacking the RNase A signal peptide. (Right). Quantification of G body formation in cells expressing plasmids with varying concentration of CuSO_4_. (B) (Left) Representative images of hypoxic Pfk2-GFP localization with varying concentrations of CuSO_4_ in cells with a vector control or plasmid expressing Pfk2-RNase A or Pfk2-RNase A^H12A^, a variant with a mutation inhibiting RNase activity. (Right) Quantification of G body formation in cells expressing plasmids with varying concentration of CuSO_4_. (A-B) For all graphs, data represent mean and standard deviation of three individual experiments with N > 100 per replicate per condition. Arrows indicate G bodies. All scale bars are 5 μm.

**Figure S5:**
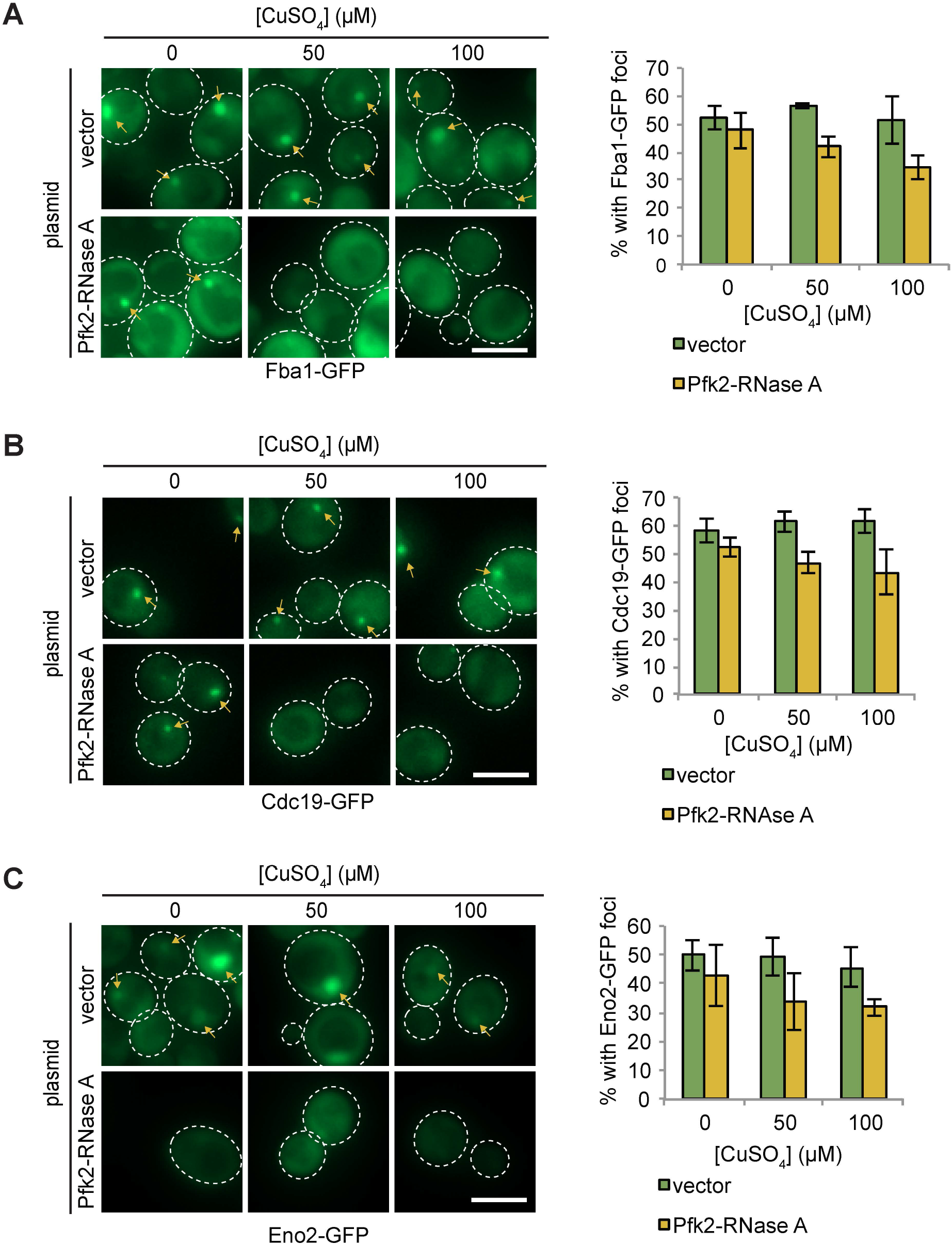
Related to Figure 3. Pfk2-RNase A inhibits punctate formation of multiple G-body markers. (A) (Left) Representative images of hypoxic Fba1-GFP localization in cells with a vector control and cells expressing Pfk2-RNase A. (Right) Quantification of punctate localization of cells. (B) (Left) Representative images of hypoxic Cdc19-GFP localization in cells with a vector control and cells expressing Pfk2-RNase A. (Right) Quantification of punctate localization of cells. (C) (Left) Representative images of hypoxic Eno2-GFP localization in cells with a vector control and cells expressing Pfk2-RNase A. (Right) Quantification of punctate localization of cells. All scale bars are 5 μm. Arrows indicate G bodies. For all graphs, data represent mean and standard deviation of three independent experiments (N > 100 cells per condition per replicate).

**Figure S6:**
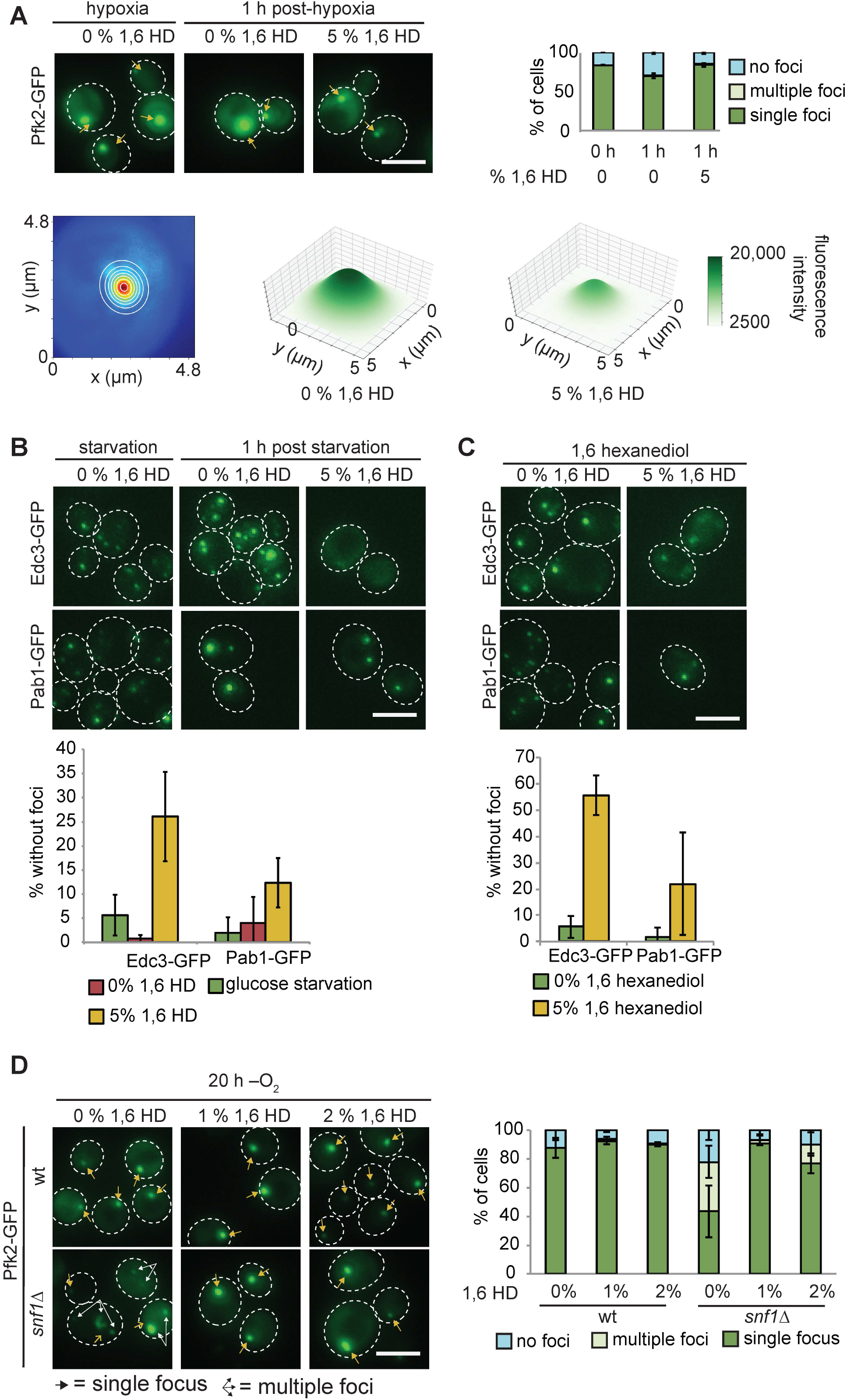
Related to Figure 6. 1,6-Hexanediol partially dissolves G bodies but does not inhibit their formation. (A) (Top) 1,6-hexanediol treatment of cells with G bodies. 5% 1,6-hexanediol was added in PBST to cells with G bodies after 20 h in hypoxia for 1 h and cells were imaged. Left: Representative images. Top Right: Quantification. (Bottom) Left: Example of 2-dimensional Gaussian distribution of Pfk2-GFP focus overlaid on real data. Blue pixels have weak fluorescence relative red pixels. Right: Average 2-dimensional Gaussian distributions fit to Pfk2-GFP foci from 5% 1,6-hexanediol treated cells following hypoxia or untreated cells. Parameters were averaged across n = 253 and 310, respectively. Notably, amplitude and standard deviations are smaller in 1,6 hexanediol treated cells (A = 18519, σ_x_ = 1.12 μm, σ_y_ = 1.12 μm and A = 12054, σ_x_ = 0.72 μm and σ_y_ = 0.74 μm respectively). (B) 1,6-hexanediol treatment of cells with Edc3-GFP (P-bodies) and Pab1-GFP (stress granules). Cells starved of glucose for 30 min were treated with 5% 1,6-hexanediol in PBST for 1 h. Left: Representative images. Right: Quantification. (C) 5% 1,6-hexanediol treatment during 30 min glucose starvation. Left: Representative images. Right: Quantification. (D) 2% 1,6-hexanediol treatment during hypoxia in wt or *snf1Δ* cells. Left: Representative images. Right: Quantification. All scale bars 5 μM. Arrows indicate G bodies. All experiments represent average +/− standard deviation of three independent experiments.

**Figure S7:**
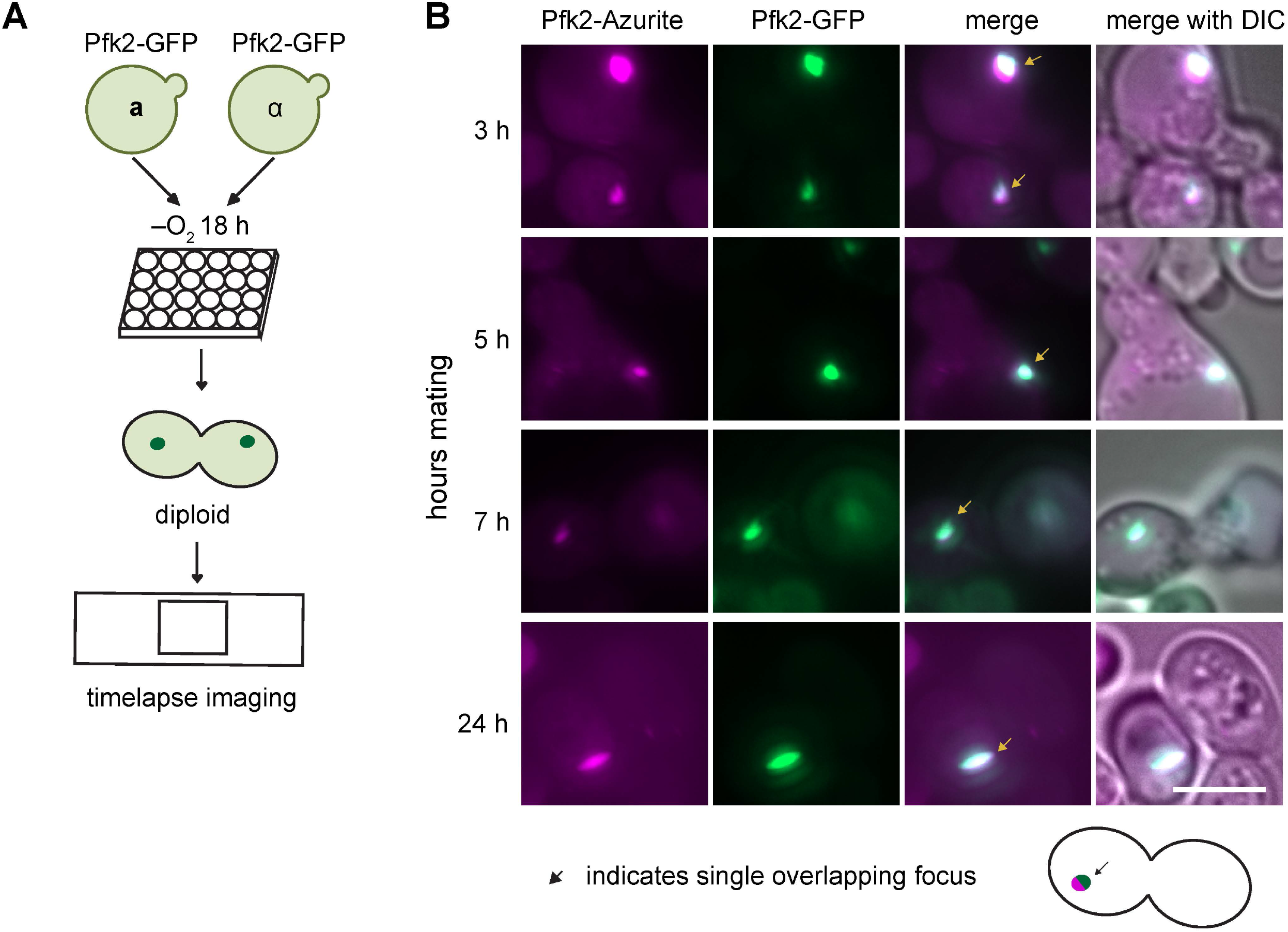
Related to Figure 6. Schematics of mating experiments. (A) Mating for kinetics measurements. a and α cells expressing Pfk2-GFP are grown together in hypoxia for 18 h. Cells are placed under a cover slip and imaged for hours with images every 2 minutes. (B) Representative images of mating cells with colocalized Pfk2-Azurite and Pfk2-GFP at each timepoint tested. Arrows indicate G bodies. Scale bar is 5 μm.

## REFERENCES

Alberti, S., Gladfelter, A., & Mittag, T. (2019). Considerations and Challenges in Studying Liquid-Liquid Phase Separation and Biomolecular Condensates. Cell, 176(3), 419–434. http://doi.org/10.1016/j.cell.2018.12.035

Amberg, D. C., Burke, D. J., & Strathern, J. N. (n.d.). Methods in Yeast Genetics: A Cold Spring Harbor Laboratory Course Manual. (D. Crotty, R. Stewer, & S. Schaefer, Eds.) (2005 ed., pp. 149–153). John Inglis.

An, S., Kumar, R., Sheets, E. D., & Benkovic, S. J. (2008). Reversible compartmentalization of de novo purine biosynthetic complexes in living cells. Science, 320(5872), 103–106. http://doi.org/10.1126/science.1152241

Baltz, A. G., Munschauer, M., Schwanhäusser, B., Vasile, A., Murakawa, Y., Schueler, M., et al. (2012). The mRNA-Bound Proteome and Its Global Occupancy Profile on Protein-Coding Transcripts. Molecular Cell, 46(5), 674–690. http://doi.org/10.1016/j.molcel.2012.05.021

Banaszak, K., Mechin, I., Obmolova, G., Oldham, M., Chang, S. H., Ruiz, T., et al. (2011). The crystal structures of eukaryotic phosphofructokinases from baker’s yeast and rabbit skeletal muscle. Journal of Molecular Biology, 407(2), 284–297. http://doi.org/10.1016/j.jmb.2011.01.019

Becerra, M., Lombardía-Ferreira, L. J., Hauser, N. C., Hoheisel, J. D., Tizon, B., & Cerdán, M. E. (2002). The yeast transcriptome in aerobic and hypoxic conditions: effects of hap1, rox1, rox3and srb10deletions. Molecular Microbiology, 43(3), 545–555. http://doi.org/10.1046/j.1365-2958.2002.02724.x

Beckmann BM, Horos R, Fischer B, Castello A, Eichelbaum K, Alleaume AM, Schwarzl T, Curk T, Foehr S, Huber W, Krijgsveld J, Hentze MW. 2015. The RNA-binding proteomes from yeast to man harbour conservedenigmRBPs. Nature Communications 6: 10127

Boeynaems, S., Alberti, S., Fawzi, N. L., Mittag, T., Polymenidou, M., Rousseau, F., et al. (2018). Protein Phase Separation: A New Phase in Cell Biology. Trends in Cell Biology, 28(6), 420–435. http://doi.org/10.1016/j.tcb.2018.02.004

Brangwynne, C. P., Eckmann, C. R., Courson, D. S., Rybarska, A., Hoege, C., Gharakhani, J., et al. (2009). Germline P granules are liquid droplets that localize by controlled dissolution/condensation. Science, 324(5935), 1729–1732. http://doi.org/10.1126/science.1172046

Brangwynne, C. P., Mitchison, T. J., & Hyman, A. A. (2011). Active liquid-like behavior of nucleoli determines their size and shape in Xenopus laevis oocytes. Proceedings of the National Academy of Sciences of the United States of America, 108(11), 4334–4339. http://doi.org/10.1073/pnas.1017150108

Buchan, J. R., Muhlrad, D., & Parker, R. (2008). P bodies promote stress granule assembly in Saccharomyces cerevisiae. The Journal of Cell Biology, 183(3), 441–455. http://doi.org/10.1083/jcb.200807043

Castello, A., Fischer, B., Eichelbaum, K., Horos, R., Beckmann, B. M., Strein, C., et al. (2012). Insights into RNA biology from an atlas of mammalian mRNA-binding proteins. Cell, 149(6), 1393–1406. http://doi.org/10.1016/j.cell.2012.04.031

Choi, H., Fermin, D., & Nesvizhskii, A. I. (2008). Significance analysis of spectral count data in label-free shotgun proteomics. Molecular & Cellular Proteomics: MCP, 7(12), 2373–2385. http://doi.org/10.1074/mcp.M800203-MCP200

Elbaum-Garfinkle, S., Kim, Y., Szczepaniak, K., Chen, C. C.-H., Eckmann, C. R., Myong, S., & Brangwynne, C. P. (2015). The disordered P granule protein LAF-1 drives phase separation into droplets with tunable viscosity and dynamics. Proceedings of the National Academy of Sciences, 112(23), 7189–7194. http://doi.org/10.1073/pnas.1504822112

Fay, M. M., & Anderson, P. J. (2018). The Role of RNA in Biological Phase Separations. Journal of Molecular Biology, 430(23), 4685–4701. http://doi.org/10.1016/j.jmb.2018.05.003

Feric, M., Vaidya, N., Harmon, T. S., Mitrea, D. M., Zhu, L., Richardson, T. M., et al. (2016). Coexisting Liquid Phases Underlie Nucleolar Subcompartments. Cell, 165(7), 1686–1697. http://doi.org/10.1016/j.cell.2016.04.047

Freeberg, M. A., Han, T., Moresco, J. J., Kong, A., Yang, Y.-C., Lu, Z. J., et al. (2013). Pervasive and dynamic protein binding sites of the mRNA transcriptome in Saccharomyces cerevisiae. Genome Biology, 14(2), R13. http://doi.org/10.1186/gb-2013-14-2-r13

Freeman Rosenzweig, E. S., Xu, B., Kuhn Cuellar, L., Martinez-Sanchez, A., Schaffer, M., Strauss, M., et al. (2017). The Eukaryotic CO2-Concentrating Organelle Is Liquid-like and Exhibits Dynamic Reorganization. Cell, 171(1), 148–162.e19. http://doi.org/10.1016/j.cell.2017.08.008

Ghaemmaghami, S., Huh, W.-K., Bower, K., Howson, R. W., Belle, A., Dephoure, N., et al. (2003). Global analysis of protein expression in yeast. Nature, 425(6959), 737–741. http://doi.org/10.1038/nature02046

Gietz, R. D., & Woods, R. A. (2002). Transformation of yeast by lithium acetate/single-stranded carrier DNA/polyethylene glycol method. Methods in Enzymology, 350, 87–96.

Hafner, M., Landthaler, M., Burger, L., Khorshid, M., Hausser, J., Berninger, P., et al. (2010). Transcriptome-wide identification of RNA-binding protein and microRNA target sites by PAR-CLIP. Cell, 141(1), 129–141. http://doi.org/10.1016/j.cell.2010.03.009

Hyman, A. A., Weber, C. A., & Jülicher, F. (2014). Liquid-Liquid Phase Separation in Biology. Annual Review of Cell and Developmental Biology, 30(1), 39–58. http://doi.org/10.1146/annurev-cellbio-100913-013325

Jain, A., & Vale, R. D. (2017). RNA phase transitions in repeat expansion disorders. Nature, 546(7657), 243–247. http://doi.org/10.1038/nature22386

Jain, S., Wheeler, J. R., Walters, R. W., Agrawal, A., Barsic, A., & Parker, R. (2016). ATPase-Modulated Stress Granules Contain a Diverse Proteome and Substructure. Cell, 164(3), 487–498. http://doi.org/10.1016/j.cell.2015.12.038

Jang, S., Nelson, J. C., Bend, E. G., Rodríguez-Laureano, L., Tueros, F. G., Cartagenova, L., et al. (2016). Glycolytic enzymes localize to synapses under energy stress to support synaptic function.

Jensen, O. N., Wilm, M., Shevchenko, A., & Mann, M. (1999). Sample preparation methods for mass spectrometric peptide mapping directly from 2-DE gels. Methods in Molecular Biology (Clifton, N.J.), 112, 513–530.

Jin, M., Fuller, G. G., Han, T., Yao, Y., Alessi, A. F., Freeberg, M. A., et al. (2017). Glycolytic Enzymes Coalesce in G Bodies under Hypoxic Stress. Cell Reports, 20(4), 895–908. http://doi.org/10.1016/j.celrep.2017.06.082

Kasari, V., Kurg, K., Margus, T., Tenson, T., & Kaldalu, N. (2010). The Escherichia coli mqsR and ygiT genes encode a new toxin-antitoxin pair. Journal of Bacteriology, 192(11), 2908–2919. http://doi.org/10.1128/JB.01266-09

Kato, M., Han, T. W., Xie, S., Shi, K., Du, X., Wu, L. C., et al. (2012). Cell-free Formation of RNA Granules: Low Complexity Sequence Domains Form Dynamic Fibers within Hydrogels. Cell, 149(4), 753–767. http://doi.org/10.1016/j.cell.2012.04.017

Kedersha, N., Stoecklin, G., Ayodele, M., Yacono, P., Lykke-Andersen, J., Fritzler, M. J., et al. (2005). Stress granules and processing bodies are dynamically linked sites of mRNP remodeling. The Journal of Cell Biology, 169(6), 871–884. http://doi.org/10.1083/jcb.200502088

Khong, A., Matheny, T., Jain, S., Mitchell, S. F., Wheeler, J. R., & Parker, R. (2017). The Stress Granule Transcriptome Reveals Principles of mRNA Accumulation in Stress Granules. Molecular Cell, 68(4), 808–820.e5. http://doi.org/10.1016/j.molcel.2017.10.015

Kohnhorst, C. L., Kyoung, M., Jeon, M., Schmitt, D. L., Kennedy, E. L., Ramirez, J., et al. (2017). Identification of a multienzyme complex for glucose metabolism in living cells. Journal of Biological Chemistry, 292(22), 9191–9203. http://doi.org/10.1074/jbc.M117.783050

Kojima, T., & Takayama, S. (2018). Membraneless Compartmentalization Facilitates Enzymatic Cascade Reactions and Reduces Substrate Inhibition. ACS Applied Materials & Interfaces, 10(38), 32782–32791. http://doi.org/10.1021/acsami.8b07573

Kroschwald, S., Maharana, S., & Simon, A. (2017). Hexanediol: a chemical probe to investigate the material properties of membrane-less compartments. Matters, 3(5), e201702000010. http://doi.org/10.19185/matters.201702000010

Kroschwald, S., Maharana, S., Mateju, D., Malinovska, L., Nüske, E., Poser, I., et al. (2015). Promiscuous interactions and protein disaggregases determine the material state of stress-inducible RNP granules. eLife, 4, e06807. http://doi.org/10.7554/eLife.06807

Langdon, E. M., & Gladfelter, A. S. (2018). A New Lens for RNA Localization: Liquid-Liquid Phase Separation. Annual Review of Microbiology, 72(1), 255–271. http://doi.org/10.1146/annurev-micro-090817-062814

Langmead, B. (2010). Aligning short sequencing reads with Bowtie. Current Protocols in Bioinformatics, Chapter 11(1), Unit 11.7–11.7.14. http://doi.org/10.1002/0471250953.bi1107s32

Li, P., Banjade, S., Cheng, H.-C., Kim, S., Chen, B., Guo, L., et al. (2012). Phase transitions in the assembly of multivalent signalling proteins. Nature, 483(7389), 336–340. http://doi.org/10.1038/nature10879

Lin, Y., Protter, D. S. W., Rosen, M. K., & Parker, R. (2015). Formation and Maturation of Phase-Separated Liquid Droplets by RNA-Binding Proteins. Molecular Cell, 60(2), 208–219. http://doi.org/10.1016/j.molcel.2015.08.018

Maharana, S., Wang, J., Papadopoulos, D. K., Richter, D., Pozniakovsky, A., Poser, I., et al. (2018). RNA buffers the phase separation behavior of prion-like RNA binding proteins. Science, 360(6391), 918–921. http://doi.org/10.1126/science.aar7366

Mann, J. R., Gleixner, A. M., Mauna, J. C., Gomes, E., DeChellis-Marks, M. R., Needham, P. G., et al. (2019). RNA Binding Antagonizes Neurotoxic Phase Transitions of TDP-43. Neuron. http://doi.org/10.1016/j.neuron.2019.01.048

Mateju, D., Franzmann, T. M., Patel, A., Kopach, A., Boczek, E. E., Maharana, S., et al. (2017). An aberrant phase transition of stress granules triggered by misfolded protein and prevented by chaperone function. The EMBO Journal, 36(12), 1669–1687. http://doi.org/10.15252/embj.201695957

Matia-González, A. M., Laing, E. E., & Gerber, A. P. (2015). Conserved mRNA-binding proteomes in eukaryotic organisms. Nature Structural & Molecular Biology, 22(12), 1027–1033. http://doi.org/10.1038/nsmb.3128

McDonald, W. H., Tabb, D. L., Sadygov, R. G., MacCoss, M. J., Venable, J., Graumann, J., et al. (2004). MS1, MS2, and SQT-three unified, compact, and easily parsed file formats for the storage of shotgun proteomic spectra and identifications. Rapid Communications in Mass Spectrometry: RCM, 18(18), 2162–2168. http://doi.org/10.1002/rcm.1603

Michels, P. A. M., Bringaud, F., Herman, M., & Hannaert, V. (2006). Metabolic functions of glycosomes in trypanosomatids. Biochimica Et Biophysica Acta, 1763(12), 1463–1477. http://doi.org/10.1016/j.bbamcr.2006.08.019

Mitrea, D. M., & Kriwacki, R. W. (2016). Phase separation in biology; functional organization of a higher order. Cell Communication and Signaling: CCS, 14(1), 1. http://doi.org/10.1186/s12964-015-0125-7

Miura, N., Shinohara, M., Tatsukami, Y., Sato, Y., Morisaka, H., Kuroda, K., & Ueda, M. (2013). Spatial Reorganization of Saccharomyces cerevisiae Enolase To Alter Carbon Metabolism under Hypoxia. Eukaryotic Cell, 12(8), 1106–1119. http://doi.org/10.1128/EC.00093-13

Molliex, A., Temirov, J., Lee, J., Coughlin, M., Kanagaraj, A. P., Kim, H. J., et al. (2015). Phase separation by low complexity domains promotes stress granule assembly and drives pathological fibrillization. Cell, 163(1), 123–133. http://doi.org/10.1016/j.cell.2015.09.015

Murakami, T., Qamar, S., Lin, J. Q., Schierle, G. S. K., Rees, E., Miyashita, A., et al. (2015). ALS/FTD Mutation-Induced Phase Transition of FUS Liquid Droplets and Reversible Hydrogels into Irreversible Hydrogels Impairs RNP Granule Function. Neuron, 88(4), 678–690. http://doi.org/10.1016/j.neuron.2015.10.030

Nagalakshmi, U., Wang, Z., Waern, K., Shou, C., Raha, D., Gerstein, M., & Snyder, M. (2008). The transcriptional landscape of the yeast genome defined by RNA sequencing. Science, 320(5881), 1344–1349. http://doi.org/10.1126/science.1158441

Ohshima, D., Arimoto-Matsuzaki, K., Tomida, T., Takekawa, M., & Ichikawa, K. (2015). Spatio-temporal Dynamics and Mechanisms of Stress Granule Assembly. PLoS Computational Biology, 11(6), e1004326. http://doi.org/10.1371/journal.pcbi.1004326

Opperdoes, F. R. (1987). Biogenesis of Glycosomes (Microbodies) in the Trypanosomatidae. In Peroxisomes in Biology and Medicine (pp. 426–435). Berlin, Heidelberg: Springer Berlin Heidelberg. http://doi.org/10.1007/978-3-642-71325-5_46

Panas, M. D., Ivanov, P., & Anderson, P. (2016). Mechanistic insights into mammalian stress granule dynamics. The Journal of Cell Biology, 215(3), 313–323. http://doi.org/10.1083/jcb.201609081

Peng, J., Elias, J. E., Thoreen, C. C., Licklider, L. J., & Gygi, S. P. (2003). Evaluation of multidimensional chromatography coupled with tandem mass spectrometry (LC/LC-MS/MS) for large-scale protein analysis: the yeast proteome. Journal of Proteome Research, 2(1), 43–50.

Protter, D. S. W., Rao, B. S., Van Treeck, B., Lin, Y., Mizoue, L., Rosen, M. K., & Parker, R. (2018). Intrinsically Disordered Regions Can Contribute Promiscuous Interactions to RNP Granule Assembly. Cell Reports, 22(6), 1401–1412. http://doi.org/10.1016/j.celrep.2018.01.036

Reimand, J., Arak, T., Adler, P., Kolberg, L., Reisberg, S., Peterson, H., & Vilo, J. (2016). g:Profiler-a web server for functional interpretation of gene lists (2016 update). Nucleic Acids Research, 44(W1), W83–9. http://doi.org/10.1093/nar/gkw199

Riback, J. A., Katanski, C. D., Kear-Scott, J. L., Pilipenko, E. V., Rojek, A. E., Sosnick, T. R., & Drummond, D. A. (2017). Stress-Triggered Phase Separation Is an Adaptive, Evolutionarily Tuned Response. Cell, 168(6), 1028–1040.e19. http://doi.org/10.1016/j.cell.2017.02.027

Saad, S., Cereghetti, G., Feng, Y., Picotti, P., Peter, M., & Dechant, R. (2017). Reversible protein aggregation is a protective mechanism to ensure cell cycle restart after stress. Nature Cell Biology, 19(10), 1202–1213. http://doi.org/10.1038/ncb3600

Scherrer, T., Mittal, N., Janga, S. C., & Gerber, A. P. (2010). A screen for RNA-binding proteins in yeast indicates dual functions for many enzymes. Plos One, 5(11), e15499. http://doi.org/10.1371/journal.pone.0015499

Schindelin, J., Arganda-Carreras, I., Frise, E., Kaynig, V., Longair, M., Pietzsch, T., et al. (2012). Fiji: an open-source platform for biological-image analysis. Nature Methods, 9(7), 676–682. http://doi.org/10.1038/nmeth.2019

Shchepachev, V., Bresson, S., Spanos, C., Petfalski, E., Fischer, L., Rappsilber, J., & Tollervey, D. (2019). Defining the RNA interactome by total RNA-associated protein purification. Molecular Systems Biology, 15(4), e8689. http://doi.org/10.15252/msb.20188689

Strater, N., Marek, S., Kuettner, E. B., Kloos, M., Keim, A., Bruser, A., et al. (2010). Crystal structure of Pichia pastoris phosphofructokinase in the T-state. http://doi.org/10.2210/pdb3opy/pdb

Tabb, D. L., McDonald, W. H., & Yates, J. R. (2002). DTASelect and Contrast: Tools for Assembling and Comparing Protein Identifications from Shotgun Proteomics. Journal of Proteome Research, 1(1), 21–26. http://doi.org/10.1021/pr015504q

Teixeira, D., Sheth, U., Valencia-Sanchez, M. A., Brengues, M., & Parker, R. (2005). Processing bodies require RNA for assembly and contain nontranslating mRNAs. Rna, 11(4), 371–382. http://doi.org/10.1261/rna.7258505

Thompson, J. E., & Raines, R. T. (1994). Value of general Acid-base catalysis to ribonuclease a. Journal of the American Chemical Society, 116(12), 5467–5468. http://doi.org/10.1021/ja00091a060

Tsai, N.-P., Ho, P.-C., & Wei, L.-N. (2008). Regulation of stress granule dynamics by Grb7 and FAK signalling pathway. The EMBO Journal, 27(5), 715–726. http://doi.org/10.1038/emboj.2008.19

Van Treeck, B., Protter, D. S. W., Matheny, T., Khong, A., Link, C. D., & Parker, R. (2018). RNA self-assembly contributes to stress granule formation and defining the stress granule transcriptome. Proceedings of the National Academy of Sciences of the United States of America, 115(11), 2734–2739. http://doi.org/10.1073/pnas.1800038115

Vasudevan, S., & Peltz, S. W. (2001). Regulated ARE-mediated mRNA decay in Saccharomyces cerevisiae. Molecular Cell, 7(6), 1191–1200.

Walters, R. W., Muhlrad, D., Garcia, J., & Parker, R. (2015). Differential effects of Ydj1 and Sis1 on Hsp70-mediated clearance of stress granules in Saccharomyces cerevisiae. Rna, 21(9), 1660–1671. http://doi.org/10.1261/rna.053116.115

Wheeler, J. R., Matheny, T., Jain, S., Abrisch, R., & Parker, R. (2016). Distinct stages in stress granule assembly and disassembly. eLife, 5, 875. http://doi.org/10.7554/eLife.18413

Wolters, D. A., Washburn, M. P., & Yates, J. R. (2001). An Automated Multidimensional Protein Identification Technology for Shotgun Proteomics. Analytical Chemistry, 73(23), 5683–5690. http://doi.org/10.1021/ac010617e

Xu, T., Park, S. K., Venable, J. D., Wohlschlegel, J. A., Diedrich, J. K., Cociorva, D., et al. (2015). ProLuCID: An improved SEQUEST-like algorithm with enhanced sensitivity and specificity. Journal of Proteomics, 129, 16–24. http://doi.org/10.1016/j.jprot.2015.07.001

Yamaguchi, Y., Park, J.-H., & Inouye, M. (2009). MqsR, a crucial regulator for quorum sensing and biofilm formation, is a GCU-specific mRNA interferase in Escherichia coli. The Journal of Biological Chemistry, 284(42), 28746–28753. http://doi.org/10.1074/jbc.M109.032904

Yassour, M., Kaplan, T., Fraser, H. B., Levin, J. Z., Pfiffner, J., Adiconis, X., et al. (2009). Ab initio construction of a eukaryotic transcriptome by massively parallel mRNA sequencing. Proceedings of the National Academy of Sciences of the United States of America, 106(9), 3264–3269. http://doi.org/10.1073/pnas.0812841106

Zhang, H., Elbaum-Garfinkle, S., Langdon, E. M., Taylor, N., Occhipinti, P., Bridges, A. A., et al. (2015). RNA Controls PolyQ Protein Phase Transitions. Molecular Cell, 60(2), 220–230. http://doi.org/10.1016/j.molcel.2015.09.017

Zhao, H., French, J. B., Fang, Y., & Benkovic, S. J. (2013). The purinosome, a multi-protein complex involved in the de novo biosynthesis of purines in humans. Chemical Communications (Cambridge, England), 49(40), 4444–4452. http://doi.org/10.1039/c3cc41437j

